# Genome-wide structural variants uncover strain-specific genes and evolutionary genomic determinants of a fungal maize pathogen

**DOI:** 10.64898/2026.10.01.755987

**Authors:** S. Becerra, P. García-Rodríguez, J. Martín, L. Rodríguez-Mónaco, S. Gutiérrez-Sánchez, F.B. Cuevas-Fernández, F. Rogério, T.R. Boufleur, J. L. Vicente-Villardón, L. Faino, R. Baroncelli, S.A. Sukno, M.R. Thon

## Abstract

Genomic *structural variants* (SVs) are important sources of genetic diversity and contribute to the adaptation and evolution of plant-pathogenic fungi. *Colletotrichum graminicola*, the causal agent of maize (*Zea mays* L.) anthracnose, causes significant yield losses and serves as an important model for genetic studies. To investigate genomic plasticity in this pathogen, long-read genome assemblies were generated for four *C. graminicola* field strains collected in France, Croatia, Brazil, and Argentina and compared with the chromosome-level reference genome from the USA. Comparative analyses revealed extensive structural variation across core chromosomes, including large-scale translocations, inversions, duplications, and insertions and deletions, whereas minichromosomes showed markedly reduced collinearity among strains. Secreted proteins and predicted effectors were located significantly closer to regions affected by SVs. Approximately 174 kb of segmental duplications lacking genes were shared between core chromosome 04 and minichromosomes 11 and 13 in the reference genome but were absent from the other field strains, revealing a strain-specific structural link between core and minichromosomes. Protein clustering identified 733 candidate strain-specific genes across the reference genome. Approximately 72% of the genes annotated on the minichromosomes were strain-specific, indicating that most of their annotated genes lacked orthologous gene models in the other strains. Of the 733 candidate strain-specific genes, 492 retained homologous genomic sequences in at least one other strain and many also showed evidence of gene disruption. This pattern is consistent with lineage-specific pseudogenization and suggests that apparent gene presence-absence variation may arise through several evolutionary processes, including gene gain, gene loss and loss of gene function. Insertions and deletions in potentially regulatory flanking regions were associated with plant-pathogen interaction genes. In addition, interpretable machine learning analyses revealed a co-occurrence of duplications and LTR retrotransposons in genomic regions associated with secreted genes. This study shows that structural variation, transposable elements, and lineage-specific pseudogenization are important features of genome evolution in *C. graminicola* and identifies associations between genomic variation and genes involved in plant-pathogen interactions.

## 2. Introduction

Ensuring long-term food security and sustainable agricultural productivity remains one of the major global challenges for modern agriculture (Cole et al., 2018; Denning, 2025). In this context, fungal plant pathogens impact crop production and sustainability worldwide. The haploid fungal plant pathogen *Colletotrichum graminicola* causes anthracnose disease in maize (*Zea mays* L.), a crop of significant global importance (Erenstein et al., 2022). This pathogen can infect all plant parts, but the most substantial economic damage is caused by stalk rot and seedling blight (Bergstrom and Nicholson, 1999; Cota et al., 2012; Palaveršić et al., 2009). In addition to its economic importance, *C. graminicola* is also an important model for studying plant-pathogen interactions (Münch et al., 2008; O’Connell et al., 2012; Sukno et al., 2008). Recently, an updated version of the *C. graminicola* M1.001 reference genome assembly was obtained using a hybrid approach combining short- and long-read sequencing technologies, supported by optical map data and cytological analysis, providing a high-resolution resource for further investigation of this pathosystem (Becerra et al., 2023; O’Connell et al., 2012; Taga et al., 2015).

Eukaryotic chromosomes can vary in size, shape, composition, and number, and may undergo changes between and within species. Structural variants such as translocations, deletions, insertions, and inversions can modify chromosome size and shape (Alkan et al., 2011; Kistler and Miao, 1992; Stukenbrock and Croll, 2014; Zolan, 1995). The genomes of phytopathogenic fungi present particularly compelling subjects for genomic investigation owing to their pronounced structural variability and genomic plasticity (Faino et al., 2016; Plissonneau et al., 2016; Tsushima et al., 2019; Zhong et al., 2020). Fungal genomes exhibit high genetic variability, which can contribute to adaptation. This genetic variability can arise through mitotic or meiotic recombination mechanisms and genomic rearrangements through ectopic recombination (Davis and Perkins, 2002; Grandaubert et al., 2019; Komluski et al., 2023; Stukenbrock and Dutheil, 2018; Wittenberg et al., 2009). A recent chromosome-level comparative genomic study in fungi revealed extensive karyotypic variation driven by chromosome fusion events and rearrangements involving centromeric regions, highlighting the contribution of large-scale genomic changes to fungal genome evolution (Coelho et al., 2024).

Members of the genus *Colletotrichum* are thought to be predominantly asexual, but for several species, the sexual states have been described under natural or controlled conditions (Barcelos et al., 2014; Souza-Paccola et al., 2003; Talhinhas and Baroncelli, 2021). In *C. graminicol*a, genetic recombination contributes to population structure and may arise through sexual or non-meiotic processes, including parasexual events (Rech et al., 2014; Rogério et al., 2025, 2023; Vaillancourt et al., 2000; Vaillancourt and Hanau, 1991). Together with structural rearrangements, these processes contribute to the high degree of genome plasticity observed in fungal plant pathogens (Stukenbrock, 2013).

In this context, fungal plant pathogens may harbor structurally distinct chromosomes, including accessory chromosomes and small chromosomes often referred to as minichromosomes, which can encode genes contributing to host infection (Bhadauria et al., 2019; Dallery et al., 2017; Pires et al., 2016). For example, in *Fusarium oxysporum,* accessory chromosomes harbor effector genes such as the *SIX* genes that are required for host-specific pathogenicity and can be horizontally transferred between strains (Ma et al., 2010). In *C. gloeosporioides*, a 2 Mb chromosome was experimentally transferred across vegetative incompatibility barriers between distinct biotypes (He et al., 1998). Minichromosomes can also harbor genes associated with virulence, and their absence in some strains can result in the loss of virulence (Covo, 2020; Kistler and Miao, 1992; Masel et al., 1993; Plaumann et al., 2018).

Certain fungi can accumulate genetic variation even during asexual reproduction without genetic recombination (De Jonge et al., 2013; Faino et al., 2016; Seidl and Thomma, 2014). Recently, the concept of clonality in fungal pathogens has been challenged by models suggesting that episodic sexual recombination generates diverse genotypes that can subsequently be maintained through clonal-like dynamics (Fayyaz et al., 2023; Hill and McMullan, 2023; Rogério et al., 2025). In this context, even infrequent recombination events introduce genetic variation that can rapidly become fixed in emerging clonal lineages. Subsequent genomic rearrangements and structural variants can further reshape genome architecture by generating variation in genomic content, including loci associated with pathogenicity, thereby contributing to divergence among lineages (Seidl and Thomma, 2014; Stukenbrock, 2013).

Transposable elements (TEs) are important contributors to genome evolution and plasticity in plant pathogens. Repetitive DNA elements such as TEs can promote genomic rearrangements through bursts of transposition, facilitating genetic changes (Badet et al., 2024; Dean et al., 2005; Faino et al., 2016; Mondo and Grigoriev, 2025; Nitta et al., 1997; Oggenfuss and Croll, 2023; Thon et al., 2006). Genomic plasticity in pathogens is also reflected in dynamic chromosomal structures such as minichromosomes, which can integrate into core chromosomes through processes including segmental duplication, thereby generating additional genomic variation (Lapalu et al., 2024; Ma et al., 2010).

Advances in sequencing technologies have enabled the resolution of genome assemblies and facilitated the detection of structural variants. Long-read sequencing enables accurate assembly of repetitive regions, allowing for a more comprehensive analysis of genomic architecture (Denti et al., 2023; Faino et al., 2015; Guo et al., 2021; Ho et al., 2020; Oggenfuss et al., 2025; Wang et al., 2025). The availability of such high-resolution genomic datasets has also encouraged the use of integrative analytical frameworks capable of modeling complex genomic features. Recent applications of machine learning approaches in fungal genomics have enabled the integration of complex genomic variables across multiple scales, from broad evolutionary comparisons to the analysis of local genomic instability (Badet et al., 2021; Fijarczyk et al., 2025).

Here, we present high-quality genome assemblies and gene annotations for four *C. graminicola* strains generated using short- and long-read sequencing technologies. We characterize intraspecific structural variation through comparative genomic analyses and investigate its genomic distribution, chromosomal organization, and association with genes involved in plant-pathogen interactions. We also use an interpretable machine learning framework to evaluate the combined associations of structural variation, transposable elements, and sequence features in relation to gene classes. This study provides new insights into the relationships between structural variation, genome plasticity, and intraspecific genome evolution in an important maize anthracnose pathogen. In contrast to pathogens whose accessory chromosomes carry host-specific effectors, we find that the *C. graminicola* minichromosomes are hypervariable and enriched in strain-specific genes yet effector-poor and that much of the apparent gene presence–absence variation reflects lineage-specific pseudogenization rather than true gene loss. These results indicate that genome plasticity in this pathogen is shaped by segmental duplications linking core and minichromosomes, together with a duplication- and LTR-associated signature near secreted genes, and they caution against relying on a single reference genome to define effector repertoires.

## 3. Methods

### 3.1. Fungal strains and DNA extraction

Four field isolates of *C. graminicola* were collected in different countries (Croatia, Argentina, Brazil, and France), providing representative samples from Europe and South America. These isolates represent three geographically isolated genetic lineages previously detected in a global collection of isolates (Rogério et al., 2023). Strain M1.001, a North American strain (O’Connell et al., 2012; Vaillancourt and Hanau, 1991), was included as the reference genome using the improved assembly CGRA01V4 (PRJNA900520) (Becerra et al., 2023).

For genome sequencing, mycelia from monosporic cultures were grown in potato dextrose broth (PDB) at 150 rpm with shaking for three days (Sanz-Martín et al., 2016). Subsequently, the mycelium was harvested by vacuum filtration, lyophilized, and stored at −80 °C. For DNA extraction, 400 mg of lyophilized mycelium was macerated into a fine powder using liquid nitrogen and a mortar and pestle and transferred to a 1.5 mL microcentrifuge tube. High molecular weight (HMW) DNA was extracted using the phenol-chloroform method described by Petersen et al. (2022) and purified with QIAGEN Genomic-Tips 20 G-1 following the manufacturer’s recommendations. DNA was quantified by fluorometry with Qubit 3.0 (Invitrogen) and the DNA Integrity Number (DIN) was determined using the Agilent Genomic DNA kit on a TapeStation 2200 system.

### 3.2. DNA sequencing and genome assembly

Strains were sequenced using a combination of short and long reads. For strain CRO-I-41 (Croatia), long reads were generated using PacBio RS *Single Molecule Real-Time* (SMRT) (Pacific Biosciences) with one Sequel II SMRT cell. Short reads were generated on a NovaSeq 6000 using a paired-end library (2 × 150 bp). Both types of data were obtained by The Centre d’Expertise et de Services (Genome Québec CES, Canada). Strains ARG-5133-7 (Argentina), BR-85925-1 (Brazil), and F-64330-2 (France) were sequenced using ONT and *DNA NanoBall Sequencing* (DNBSeq), both from BGI Genomics (Hong Kong, China) (Supplementary Table S1).

The quality of the short reads was checked with FastQC v.0.11.9. Low-quality reads and adapters were removed with Trim Galore v.0.6.4 and paired-end reads were merged with FLASH v.1.2.7 (Magoc and Salzberg, 2011). LongQC v.1.2.0.b (Fukasawa et al., 2020) was used to verify the quality of long reads. Porechop v.0.2.4 (https://github.com/rrwick/Porechop) was used to remove the adapters from the ONT raw data. Filtlong v.0.2.1 (https://github.com/rrwick/Filtlong) was used with the parameter `--min_length 20000` to filter the raw long reads by quality and length. The *de novo* genome assembly of CRO-I-41 was obtained using Canu v.2.1.1 from PacBio data. For the remaining strains, the minimap2-miniasm pipeline was used, generating overlaps among ONT long reads using Minimap2 v.2.24 (Li, 2018). Miniasm v.0.3 (Li, 2016) was used to generate the contigs and Racon v.1.3.1 was used to polish the assemblies with the long-read data, followed by polishing using Pilon v.1.23 (Walker et al., 2014) with the short reads.

The sequence assemblies were manually verified using D-GENIES (Cabanettes and Klopp, 2018), pyGenomeViz (https://github.com/moshi4/pyGenomeViz) with the MUMmer option, and SyMAP v.5.1.0 (Soderlund et al., 2006) to visualize the syntenic contigs. Contigs were reorganized and oriented based on similarity to the M1.001 assembly. LongStitch (Coombe et al., 2021) was used for scaffolding with the complete long-read dataset. Finally, three iterations of Pilon v.1.23 (Walker et al., 2014) were used to obtain the final version of the assembly for each strain. These genome assemblies were tested for genome completeness using BUSCO v.5.2.2 (Manni et al., 2021) with the `sordariomycetes_odb10` lineage dataset. The genome assembly statistics were calculated using QUAST v.5.2.0 (Gurevich et al., 2013).

### 3.3. Genome and functional annotation

Gene annotation was carried out following the protocol described by Becerra et al. (2023), utilizing their species-specific training database for *C. graminicola*. Proteins containing a signal peptide, as determined by SignalP v.5.0.b (Almagro Armenteros et al., 2019), were further analyzed using EffectorP v.3.0.0 (Sperschneider and Dodds, 2022). Carbohydrate-active enzymes (CAZymes) were predicted using run_dbCAN v.4 (Zheng et al., 2023) with the parameter `--use_signalP=TRUE` to incorporate signal peptide prediction into the analysis. The PHI-base database (Urban et al., 2020) was used to identify homologous proteins related to pathogen-host interactions using blast, considering hits with an *E*-value ≤ 1 × 10^-5^ as significant.

### 3.4. Phylogenetic relationships between the strains

A phylogenomic tree was constructed using the four genomes sequenced in this study, plus the reference genome CGRA01V4 (Becerra et al., 2023), and *C. tofieldiae* (NCBI: GCA_022836555.1) as an outgroup. OrthoFinder v.2.5.4 (Emms and Kelly, 2019) was used to identify single-copy orthogroups, which were then aligned using MAFFT v.7.453 (Katoh and Standley, 2013). Poorly aligned regions were removed using Gblocks v.0.91b (Talavera and Castresana, 2007). A phylogenetic tree was built using FastTree v.2.1.11 (double-precision build, no SSE3) (Price et al., 2010).

### 3.5. Identification of structural variants

Conservation of synteny was determined with pyGenomeViz v.0.4.3 using MUMmer, with a threshold of 70% identity (Shimoyama, 2022). SyRI v.1.6.3 (Goel et al., 2019) was used to identify and classify structural rearrangements in pairwise comparisons of each genome assembly against the CGRA01V4 reference genome. Plotsr v.0.4.1 (Goel and Schneeberger, 2022) was used to visualize the results from SyRI. The coverage of each structural variant type was determined with `bedtools coverage`. Distances from genes in each category to the nearest structural variant were calculated using `bedtools closest` with the parameters `-d -t first`. The statistical significance of the observed distances between each gene category and structural variants was assessed using regioneR (Gel et al., 2016). This statistical analysis was performed with 10,000 permutations, following the methodology of Dallery et al. (2017). Regions identified as syntenic to the reference strain were compiled. The percentage of overlapping base pairs between the syntenic regions and the reference strain was calculated for each chromosome, using `bedtools coverage` in 10 kb windows. An additional chromosome-level analysis was performed to determine collinearity between sequences and gene content using MCScanX (Wang et al., 2012).

### 3.6. Identification of segmental duplications, strain-specific regions/genes

Segmental duplications (SD) in the reference genome were identified using SDDetector following the protocol described by Dallery et al. (2017). These regions were extracted and submitted to megablast to confirm their presence in the assembled strains. Subsequently, these regions were manually verified using blastn to confirm their uniqueness and identify them as strain-specific regions. Protein clustering was performed using blastclust from blast v.2.2.19 with a sequence identity threshold of 80% (`-S 80`) to identify candidate strain-specific genes. Proteins unique to each strain were subsequently compared against the proteomes of the remaining strains using blastp. Proteins showing 100% sequence identity and 100% query coverage were removed, and the remaining candidates were manually verified by blastn to eliminate false positives, yielding the final set of genes lacking orthologous annotated proteins in the other strains.

SnpEff (Cingolani et al., 2012) was used in Galaxy Version 4.3+T.galaxy2 to identify genes carrying high-impact variants. This analysis used the VCF files generated by SyRI for pairwise comparisons of CRO-I-41, ARG-5133-7, F-64330-2, and BR-85925-1 against CGRA01V4. Variant observations were summarized separately for each comparison. Variant density was calculated by dividing the number of reference base pairs represented in each comparison by the corresponding number of variant observations and was expressed as one variant observation per N bp. Minichromosome variants were available only for CRO-I-41 and ARG-5133-7. SnpEff impact categories were summarized as predicted effects and kept separate from variant observations because a single variant can generate more than one predicted effect. A gene was classified as affected by a predicted high-impact effect when its gene-level SnpEff table contained at least one high-impact annotation. Finally, candidate strain-specific genes were evaluated using genomic DNA comparisons to distinguish apparently absent loci from loci retaining homologous genomic sequences. Predicted high-impact effects were then examined separately as evidence of potential gene disruption. Candidate strain-specific genes were analyzed with InterProScan v.5.55-88.0 (Jones et al., 2014). Those with known functions underwent manual curation to group them into functional categories. To determine the significance of the distance between candidate strain-specific genes and structural variants, the regioneR package was used with the *permTest* function and 10,000 permutations.

### 3.7. Maize anthracnose virulence assays

Anthracnose leaf blight assays were performed on maize plants of line Mo940 at the V3 developmental stage following the procedure described by Vargas et al. (2012). Maize plants were placed horizontally on moist trays and inoculated with conidial suspensions of *C. graminicola* strains ARG-5133-7, BR-85925-1, CRO-I-41, F-64330-2, and M1.001, represented by the CGRA01V4 reference assembly. Lesion development was evaluated and measured at four days post-inoculation. For each strain, four independent biological replicates were evaluated, with three leaves per replicate and 20 inoculation sites per leaf. Differences in virulence among strains were evaluated using a linear mixed-effects model. Individual necrotic lesion area was included as the response variable, strain as a fixed effect, and leaf identity as a random intercept to account for the non-independence of multiple inoculation sites measured on the same leaf. The model was fitted in R using the lme4 package (Bates et al., 2015) and the significance of the fixed effect was evaluated using *lmerTest* (Kuznetsova et al., 2017). *P*-values from pairwise comparisons were adjusted using Tukey’s method, and differences were considered statistically significant at *P* < 0.05.

### 3.8. Machine learning-based feature evaluation of genes involved in plant-pathogen interactions

#### Data processing and preparation

Preliminary analyses were conducted to evaluate variable overlap across three protein categories: non-secreted, secreted proteins without a significant PHI-base match, and Secreted-PHI-base proteins, defined as secreted proteins with a blast hit against plant-pathogen entries in PHI-base (*E*-value ≤ 1 × 10^-5^). These analyses compared variables related to nucleotide composition, structural variants (SVs), and transposable elements (TEs) among the three protein categories across the gene region and the upstream and downstream flanking regions. The initial objective was to clearly separate secreted proteins from those with a significant match to PHI-base entries (Secreted-PHI-base). However, not all proteins potentially involved in pathogenicity are represented in PHI-base, and the absence of a significant blast match does not mean that a secreted protein is not involved in virulence or host interactions. Therefore, the two secreted categories were combined for the subsequent analyses.

Consequently, the analysis was approached as a binary classification task, categorizing secreted genes as the positive class and non-secreted genes as the negative class. SVs were identified by comparing the four strains used in this study with the reference strain (CGRA01V4). From these SVs, two sets of values were generated: a binary matrix (*has_*) indicating the presence (1) or absence (0) of each SV in the genomes, and a smoothed proportion (*prop_smooth*) representing the occurrence of each SV across the strains included in the analysis. The proportion was calculated considering the number of strains in which the SV was observed relative to the number of strains available for each comparison. This approach accounts for differences in strain availability among comparisons.

Additionally, TEs were classified into major classes and subclasses using HMM profiles from the Dfam database v.3.8 (Storer et al., 2021). The results were manually curated using Dfam targets and integrated with RepeatMasker data to ensure accurate classification. Genomic features such as GC content and nucleotide proportions were included. Variables were analyzed across three defined genomic regions: a *gene* region corresponding to the 15,118 annotated genes in the reference genome, and two additional 10,000 bp flanking regions, one *upstream* and one *downstream* of each gene.

To comprehensively evaluate feature importance, three feature-block configurations (Feature Blocks) were created. The complete mapping of individual variables to each feature block is provided in Supplementary Table S2, and the biological classification and metadata of all features are provided in Supplementary Table S3. These sets were designed by grouping or merging specific variables to address distinct aspects of SVs, TEs, and general genomic features, thereby facilitating a more accurate assessment of their biological relevance. All variables were derived from the master dataset, which is described in Supplementary Table S4.

<u>Feature Block 1</u> consisted of the full feature set, including each SV type in each gene-associated region, TE presence variables for individual classification, and sequence composition features such as GC content and nucleotide proportions. This block was designed as a baseline configuration to evaluate the contributions of individual genomic features. An additional version of Feature Block 1 excluding GC and nucleotide-composition variables was tested during model development but was not included in the subsequent analyses.

<u>Feature Block 2</u> grouped variables related exclusively to *complete*-region features, retaining SV density variables summarized across the full gene-associated region and TE presence variables grouped by TE classification. This block enabled the evaluation of the cumulative effects of SVs and TEs across the combined *upstream*, *gene*, and *downstream* regions.

<u>Feature Block 3</u> was a reduced and refined configuration designed to limit redundancy and improve interpretability. TE variables were aggregated into major biological categories (for example, LTR and Non-LTR retrotransposons), and a reduced set of complete-region SV variables was retained rather than separate *upstream*, *gene*, and *downstream* representations.

Each feature was evaluated individually and in combination across these *Feature Blocks*. This hierarchical feature-block design enabled a systematic exploration of the genomic determinants that contribute to the classification of genes potentially involved in plant-pathogen interactions, while maintaining consistency across modeling strategies.

#### Model application and evaluation

The evaluation of the importance of the variables, grouped into three sets called “Feature Blocks”, was performed using the *eXtreme Gradient Boosting* (XGBoost) algorithm (Chen and Guestrin, 2016), known for its ability to capture complex relationships in biological and environmental data (Chi et al., 2022; De Oliveira Aparecido et al., 2024; Sia et al., 2025). To enhance the interpretability and robustness of the analysis, a systematic multi-stage methodology was applied. Data splitting was performed using `GroupShuffleSplit`, ensuring that genes from the same genomic region were not shared between the training and test sets to avoid information leakage. Genes were grouped into fixed-size genomic bins based on chromosome and start position. Because the largest chromosome was 7.56 Mb, the 10-Mb bin size resulted in each bin encompassing one complete chromosome-level contig, thereby ensuring reduced spatial overlap between the training and testing partitions. Hyperparameter optimization was performed for each Feature Block using Optuna v.4.1.0 (Akiba et al., 2019), with the objective of maximizing the F1-score. Model performance was assessed using cross-validation, applying group-aware splitting when grouping information was available. Class imbalance was addressed within the model by optimizing the `scale_pos_weight` parameter rather than applying external resampling techniques. This approach allowed the model to account for imbalance while preserving the original data distribution. Rather than applying a fixed classification threshold, the decision threshold was optimized using the precision–recall curve, selecting the threshold that maximized the F1-score during cross-validation. The mean optimal threshold across folds was then applied to the held-out test set. Model performance was evaluated using F1-score, precision, recall, ROC-AUC, and average precision.

Feature relevance was assessed using two complementary approaches derived from the previously trained XGBoost model: Gain-based importance (Chen and Guestrin, 2016) and *Shapley additive explanations* v.0.47.0 analysis (SHAP) (Lundberg and Lee, 2017) which were used to quantify both global feature importance and sample-level contributions. SHAP summary plots and dependence plots were generated to explore the direction and magnitude of feature effects. For the XGBoost waterfall plot, SHAP values were reported on the model’s raw-margin (log-odds) scale, and additivity between the expected value, feature contributions, and model output was verified for the plotted gene. Due to correlation among genomic features and the non-linear nature of model responses, partial dependence plots (PDPs) and individual conditional expectation (ICE) curves (Goldstein et al., 2015) were interpreted qualitatively. For the main interpretability configuration, PDPs and ICE curves were computed using training data to describe model-learned trends for the most influential SV and TE features.

The robustness of the results was assessed by repeating `GroupShuffleSplit` with 30 prespecified random seeds (1000-1029) using the genomic bins described above as groups and the hyperparameters obtained from the reference run. For each seed, SHAP importance was calculated across the full dataset using the same reference dataset, and agreement among feature rankings was evaluated using tie-corrected Kendall’s *W*. The relationship between univariate Spearman correlations and mean absolute SHAP importance was evaluated using Kendall’s *tau*. Because *INDEL_complete* showed high SHAP importance but a weak univariate association, its relationship with the model output was further examined using SHAP dependence plots, binned SHAP values and approximate feature interactions.

## 4. Results

### 4.1. Near chromosome-level genome assemblies and gene annotations for four strains of *C. graminicola*

Genome assemblies were generated for four field isolates of *C. graminicola*: CRO-I-41 (Croatia), ARG-5133-7 (Argentina), F-64330-2 (France), and BR-85925-1 (Brazil). Based on the population structure described by Rogério et al. (2023), CRO-I-41, ARG-5133-7, and F-64330-2 belong to the European lineage, whereas BR-85925-1 belongs to the Brazilian lineage. The previously published North American strain M1.001 was included as the reference genome, using the improved assembly CGRA01V4 (PRJNA900520) (Becerra et al., 2023). Together, the five strains represent the three major genetic lineages identified in the global collection analyzed by Rogério et al. (2023).

For CRO-I-41 and ARG-5133-7, chromosome-level genomic assemblies were obtained, while for BR-85925-1 and F-64330-2, near chromosome-level assemblies were obtained. The assemblies varied in size, with ARG-5133-7 being the smallest and BR-85925-1 the largest (Table 1). The strains CRO-I-41 and ARG-5133-7 possessed 12 chromosomes, each with two minichromosomes, less than 1.5 Mb in length. In contrast, the strains BR-85925-1 and F-64330-2 had 18 sequences, eight of which are less than 1.5 Mb. Based on species-specific definitions of chromosome size, the chromosomes were categorized into two distinct groups. The first group, comprising chromosomes 1 to 10, is referred to here as the core chromosomes. The second group comprises the minichromosomes, which are smaller contigs and are numbered from 11 onward. The number of core chromosomes was constant across the strains analyzed, whereas the number of putative minichromosomes varied.

**Table 1.** Summary of sequencing statistics and genome features of *C. graminicola* strains.

| Description | CGRA01V4* | CRO-I-41 | ARG-5133-7 | BR-85925-1 | F-64330-2 |
| --- | --- | --- | --- | --- | --- |
| Total Length (bp) | 57,426,633 | 57,709,034 | 56,382,330 | 57,756,662 | 56,580,618 |
| Number of sequences | 13 | 12 | 12 | 18 | 18 |
| N50 (bp) | 6,444,216 | 6,699,630 | 6,409,883 | 6,441,030 | 6,503,001 |
| L50 | 5 | 4 | 4 | 5 | 4 |
| GC (%) | 45.98 | 45.89 | 46.05 | 45.88 | 46.28 |
| Ns per 100 kbp | 0.32 | 0.00 | 0.20 | 0.19 | 0.35 |
| Complete BUSCO (%) | C:98.2 [S:97.9, D:0.3] | C:98.2 [S:98.0, D:0.2] | C:97.3 [S:97.1, D:0.2] | C:97.7 [S:97.5, D:0.2] | C:98.1 [S:97.9, D:0.2] |
| Number of predicted genes | 15,118 | 15,010 | 14,743 | 14,959 | 14,995 |
| Genes with RNA-Seq evidence (%) | 92.00 | 91.68 | 91.60 | 91.49 | 91.69 |
| Number of predicted CAZymes | 1,029 | 1,074 | 1,052 | 1,065 | 1,052 |
| Number of predicted secreted proteins (SignalP) | 1,474 | 1,459 | 1,423 | 1,451 | 1,439 |
| Number of predicted effector proteins (EffectorP) | 521 | 510 | 496 | 506 | 503 |
| Repetitive DNA elements (%) | 26.00 | 27.72 | 26.88 | 27.40 | 26.37 |
| Predicted RIP sites (%) | 27.41 | 27.99 | 27.63 | 28.29 | 26.57 |

\*Reference metrics from Becerra et al. (2023) are shown for comparison between *C. graminicola* strains.

Analysis of the gene models with BUSCO showed completeness values above 97% (Table 1). The number of predicted genes was similar to the reference genome, ranging from 14,743 in ARG-5133-7 to 15,010 in CRO-I-41. More than 91% of predicted genes were supported by RNA-Seq evidence. Repetitive DNA elements constituted more than 26% of the genomes.

Overall, only minor differences in the number of predicted CAZymes, secreted proteins and predicted effectors were observed among strains. The CGRA01V4 genome contains the highest number of secreted proteins, totaling 1,474, while ARG-5133-7 has the lowest, at 1,423. The number of predicted effector proteins identified in each strain ranged from 496 to 521 (Table 1).

### 4.2. Most core chromosomes are highly syntenic and minichromosomes are structurally variable

Synteny analysis of the chromosome-level genome assemblies of CGRA01V4, ARG-5133-7, and CRO-I-41 revealed large blocks of collinearity among the core chromosomes. However, several substantial intrachromosomal translocations and inversions were found in chromosomes chr04 and chr09 (Figure 1A). Consistent with synteny plots, SyRI identified 788 translocations and 47 inversions in ARG-5133-7, compared with 675 translocations and 43 inversions in CRO-I-41. The strain from Argentina also exhibited a higher proportion of structurally variable regions, including more extensive non-aligned and highly diverged segments than CGRA01V4. Despite these rearrangements, most of the genome remained syntenic, especially across the core chromosomes.

**Figure 1.**
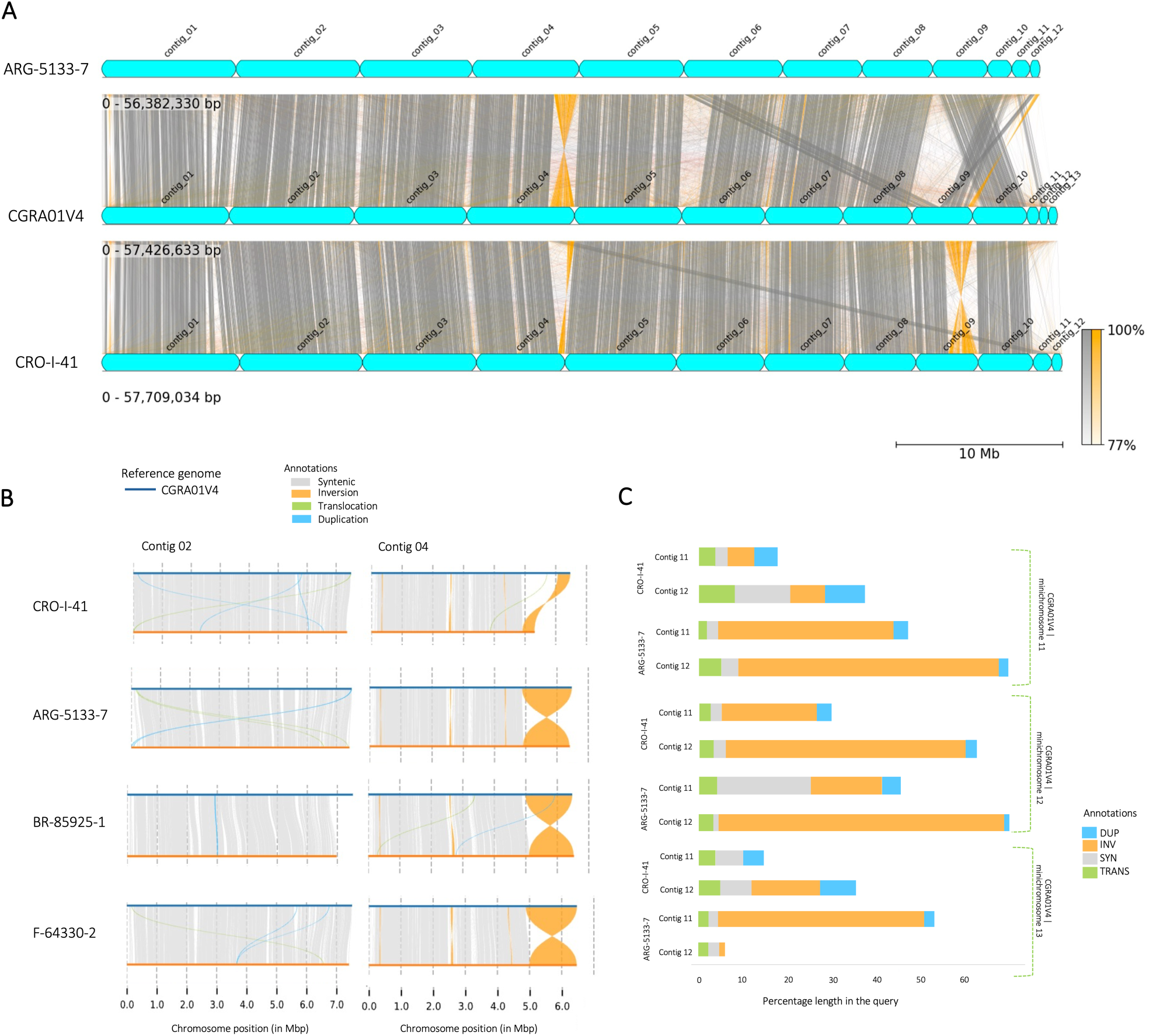
Chromosome-level synteny and structural rearrangements in *Colletotrichum graminicola*. A) NUCmer alignment visualized by pyGenomeViz, including the complete genomes for ARG-5133-7 and CRO-I-41 compared with CGRA01V4. Each genome assembly is represented by a blue box showing the contigs. Gray lines show regions of synteny between each comparison, while orange lines indicate inversions. B) Plotsr using SyRI output for four strains of *C. graminicola* with respect to the reference genome (CGRA01V4), showing Contig 02 with a high level of synteny and Contig 04 with a large inversion. C) Histogram showing the total percentage of large-scale structural variants (DUP = duplication, INV = inversion, TRANS = translocation) and syntenic (SYN) regions of the minichromosomes with respect to the three minichromosomes of the reference strain CGRA01V4, applied for the Argentine and Croatian strains in each pairwise comparison. *In the reference genome, contig = chromosome*.

The synteny analysis also revealed both synteny conservation and structural rearrangements among the minichromosomes. For instance, chr11 of ARG-5133-7 showed regions of synteny with minichromosome chr12 of CGRA01V4, whereas chr12 of ARG-5133-7 was syntenic with core chr09 of CGRA01V4, which does not display typical minichromosome characteristics (Figure 1A). Chr11 of ARG-5133-7 and chr12 of CRO-I-41 shared features with three CGRA01V4 minichromosomes, including sizes below 1.5 Mb, low gene density, and enrichment of repetitive elements. These two sequences were also evaluated for similarity to the characteristics of the three minichromosomes of CGRA01V4. Chr12 from CRO-I-41 and chr11 from ARG-5133-7 had high repetitive element content (63% and 71%, respectively) and low gene density (3% and 7%, respectively), as detailed in Supplementary Table S5. Furthermore, they exhibited the highest synteny with the minichromosomes of CGRA01V4 (Figure 1A, C). In contrast, chr11 from CRO-I-41 and chr12 from ARG-5133-7 showed similar characteristics to core chromosomes.

A secondary synteny analysis was performed by comparing the CGRA01V4 genome with each of the four previously described genomes using SyRI (Supplementary Table S6). Among the core chromosomes with fewer structural variants, chr02, chr05, and chr07 showed synteny exceeding 93%, with the majority of large-scale structural variants being duplications (DUP) and translocations (TRANS). Additionally, chr01, chr03, and chr08 showed synteny ranging from 81% to 88%, predominantly with DUP and TRANS events. Chr04 and chr09 showed extensive structural variation across the four strains, as did chr06, especially in relation to the strain from Brazil. For chr10, synteny was 91% in comparisons with CRO-I-41 and F-64330-2, but lower in comparisons with ARG-5133-7 and BR-85925-1 (Supplementary Figure S1). Among the core chromosomes, chr04 showed comparatively lower synteny (Figure 1B), which prompted us to further assess collinearity at the gene level.Despite the lower sequence-level synteny, gene collinearity on chr04 exceeded 98%.

The minichromosomes predominantly contain non-syntenic regions. The minichromosomes have a lower percentage of synteny (32%) than the core chromosomes, indicating greater structural variation. Minichromosomes chr11 and chr13 have 18% and 25% synteny, respectively, reflecting the low degree of collinearity among them (Figure 1C, Supplementary Figure S1). Across the four pairwise comparisons, 3,349,757 predicted effects were identified. Of these, 2,764,370 (82.52%) were intergenic. At the gene level, 1,505 of the 15,118 annotated genes (9.96%) were associated with at least one predicted high-impact effect. Among these genes, 131 were predicted to encode secreted proteins. Among these 1,505 genes, 73% had no known function, while the remaining genes had multiple functions, such as MFS transporters, cytochrome P450s and fungal-specific transcription factors (Supplementary Table S7).

Pairwise SyRI comparisons against CGRA01V4 identified 767,014 variant observations in CRO-I-41, 763,566 in ARG-5133-7, 702,281 in F-64330-2, and 680,227 in BR-85925-1. Variant densities ranged from one variant observation per 74.9 bp to one per 79.2 bp. Across the four comparisons, a total of 2,913,088 variant observations were identified. This total represents the sum of the four pairwise comparisons rather than the number of unique variant loci. Among large-scale structural rearrangements, TRANS, DUPs and INVs were identified, and the genome exhibited a high incidence of local INDELs (Figure 1C).

In the core chromosomes, no consistent pattern of structural variation was observed across strains. In contrast, the minichromosomes exhibited reduced synteny and a higher incidence of structural variants, predominantly INVs (Figure 1C). INDELs were identified throughout the genome, including both core chromosomes and minichromosomes and were the most prevalent variant type in all four strains compared with the reference genome.

### 4.3. Segmental duplications link core chromosome 04 with minichromosomes 11 and 13

The CGRA01V4 reference genome exhibits three segmental duplication blocks totaling approximately 174 kb, which also constitute the strain-specific regions, involving chr04, chr11 and chr13 (Figure 2A). These regions are enriched in repetitive elements, repeat-induced point mutations (RIPs), and structural variants, while showing depletion of genes (Figure 2B). These duplicated regions connect core chr04 with minichromosomes 11 and 13 and are enriched in repetitive elements and structural variation (Figure 2B). Although segmental duplications were also detected in the other strains (Supplementary Figure S2), the specific ∼174 kb duplicated regions linking chr04 with chr11 and chr13 were unique to CGRA01V4.

**Figure 2.**
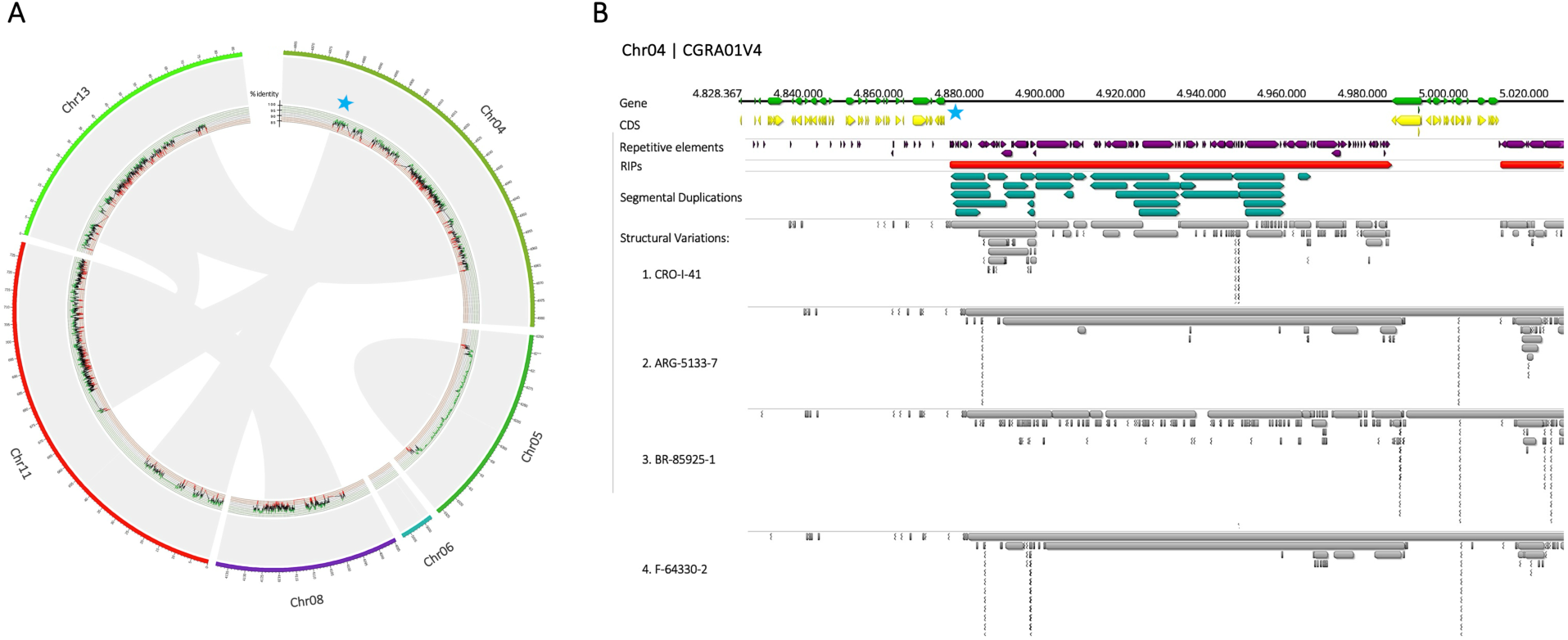
Segmental duplications linking core and minichromosomes in *Colletotrichum graminicola*. A) Circos plot of the segmental duplications (SD) in the reference genome CGRA01V4. Only regions with SD are shown. Gray arcs show the region affected by segmental duplications. Red and green lines show the coverage of repetitive elements. B) A region of chr04 showing the SD strain-specific region shared between the core and minichromosomes. This region contains an abundance of repetitive elements and structural variants compared to the other strains. Blue lines show the SD strain-specific linking core chr04 with two minichromosomes (chr11 and chr13). The blue star marks the approximate corresponding position in panels A and B.

### 4.4. Candidate strain-specific genes are located in highly dynamic regions

Candidate strain-specific genes were initially identified based on the absence of orthologous proteins in the other annotated proteomes. However, for the reference genome the subsequent analyses revealed that a proportion of these loci retained homologous genomic DNA sequences in other strains despite lacking corresponding gene models. Therefore, the strain-specific classification used here refers to the absence of an orthologous annotated gene rather than necessarily to the complete absence of the corresponding genomic locus. Among the 733 candidate strain-specific genes identified in the reference genome, 492 (67%) retained homologous genomic sequences in at least one other strain but lacked a corresponding gene model. Of these 492 loci, 333 were also affected by at least one predicted high-impact effect, supporting lineage-specific pseudogenization. Thus, some genes classified as strain-specific may reflect differences in gene integrity among strains rather than complete gain or loss of the corresponding genomic sequence.

Most core chromosomes showed a low percentage of candidate strain-specific genes, ranging from 2% to 6%, in contrast to the high percentages found in minichromosomes 11, 12, and 13. On minichromosome 12, 28% of the annotated genes were classified as strain-specific, compared with 93% on minichromosome 11 and 100% on minichromosome 13. Interestingly, among the 121 genes annotated in the three minichromosomes of the reference genome, 87 were classified as candidate strain-specific. Although these genes are candidate strain-specific, predicted secreted proteins have not been described in these minichromosomes in the reference genome. In addition, the association between different gene groups and structural variants was examined by measuring the distance of each gene to its nearest variant. This analysis revealed that predicted effector genes, genes encoding secreted proteins, and candidate strain-specific genes were located significantly closer to structural variants than expected compared to the rest of the genes (10,000 permutations; *P* ≤ 0.0001) (Figure 3A). An additional analysis tested whether genes in each category were more likely to overlap a structural variant. Predicted effector genes, genes encoding secreted proteins, and candidate strain-specific genes were more likely to overlap a structural variant than the remaining genes in the genome (Figure 3B).

**Figure 3.**
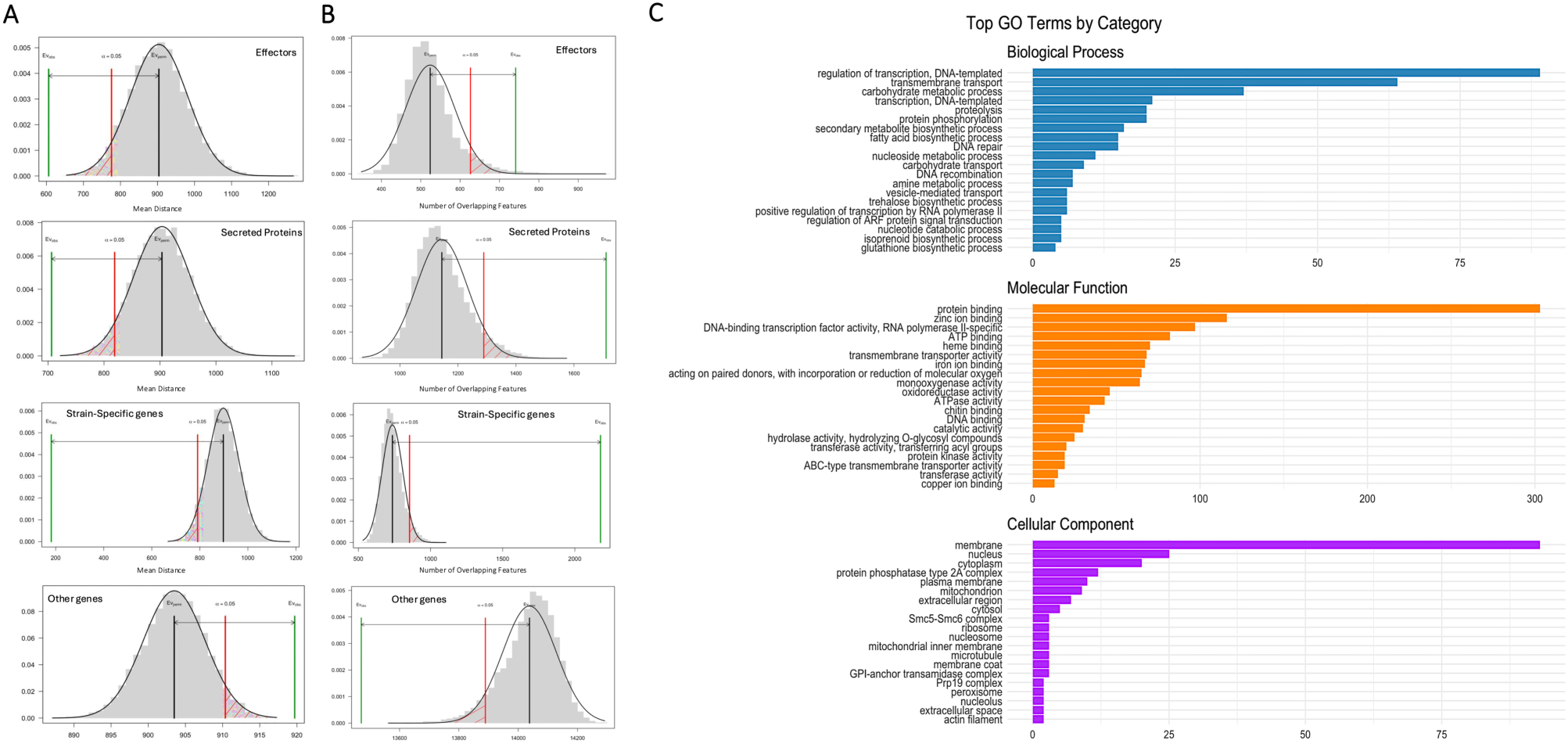
Enrichment of structural variation near candidate strain-specific genes in *Colletotrichum graminicola*. A) Distance (bp) from genes in each category to the nearest structural variant: putative effector genes (n = 521), secreted proteins (n = 1,474), other genes (n = 13,644), and candidate strain-specific genes (n = 733). Statistical significance of the distance between each gene category and structural variants was assessed using the regioneR package with 10,000 permutations. B) Number of genes that overlap SVs. C) Most frequent Gene Ontology (GO) terms associated with candidate strain-specific genes, grouped into biological process, molecular function, and cellular component categories. Bar length indicates the frequency of each GO term.

ARG-5133-7 had 125 candidate strain-specific genes, F-64330-2 had 148, CRO-I-41 had 200, BR-85925-1 had 366 and CGRA01V4 had 733 genes. The distribution broadly reflected the phylogenomic relationships, with the closely related ARG-5133-7 and F-64330-2 strains showing similarly low numbers of candidate strain-specific genes, whereas BR-85925-1 and CGRA01V4 showed higher numbers (Figure 4A, B). The candidate strain-specific genes in CGRA01V4 were analyzed for predicted high-impact effects in comparisons with the four other strains. These effects include frameshifts or changes affecting stop codons, which are predicted to disrupt gene function. Notably, 457 of the 733 candidate strain-specific genes (62%) were affected by at least one predicted high-impact effect in at least one of the four other strains. By comparison, only 1,048 of the remaining 14,385 genes (7.3%) were affected by at least one predicted high-impact effect, indicating the higher prevalence of predicted high-impact effects among candidate strain-specific genes.

**Figure 4.**
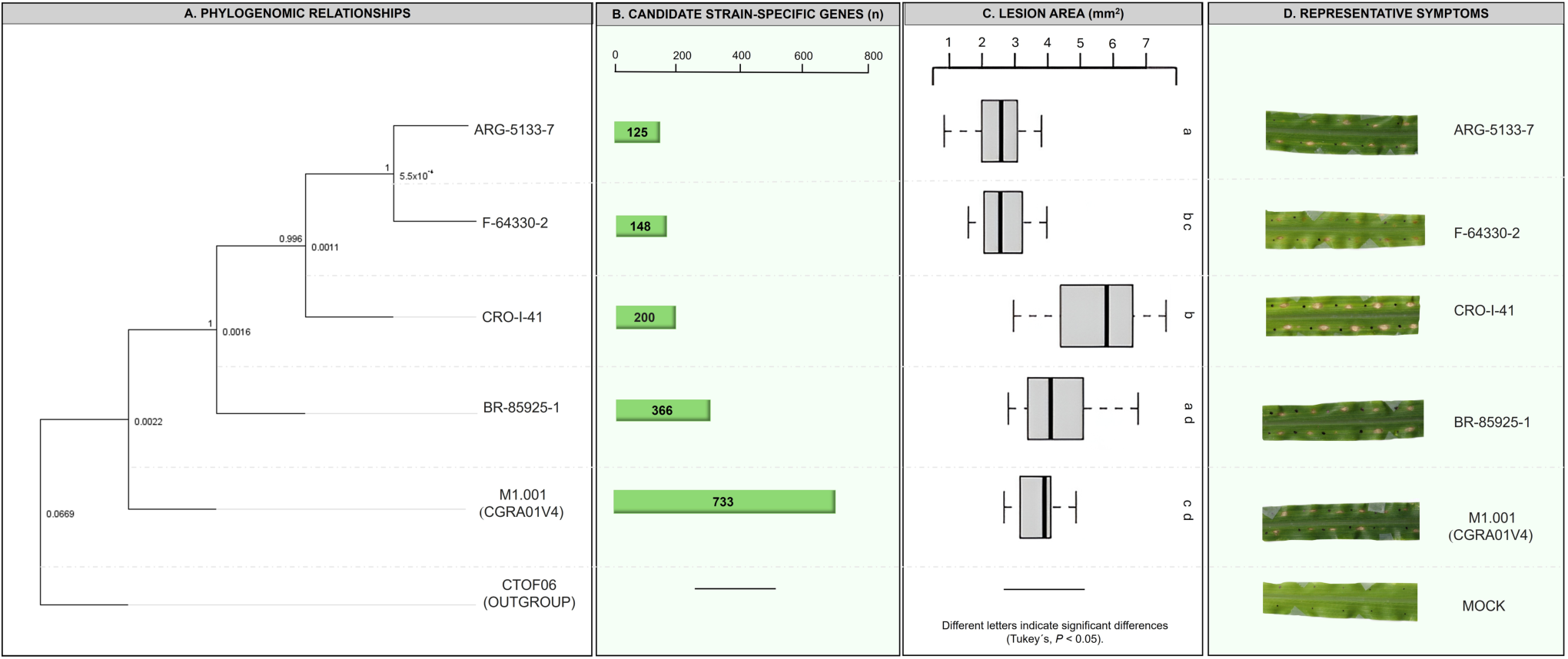
Phylogenomic relationships, candidate strain-specific genes, and virulence of *Colletotrichum graminicola* strains. A) Phylogenomic tree constructed from single-copy orthogroups identified with OrthoFinder, aligned with MAFFT, filtered with Gblocks, and inferred with FastTree. *Colletotrichum tofieldiae* CTOF06 (GCA_022836555.1) was included as the outgroup. B) Number of candidate strain-specific genes identified in each strain. C) Necrotic lesion area on maize at four days post inoculation. Different letters indicate significant differences among strains based on Tukey-adjusted pairwise comparisons of estimated marginal means from the linear mixed-effects model (*P* < 0.05). D) Representative anthracnose symptoms produced by each strain.

Functional annotation of candidate strain-specific genes revealed differences in the representation of predicted functional categories among strains. However, the majority of these genes have unknown functions. Gene Ontology analysis of candidate strain-specific genes identified frequent terms related to transcriptional regulation, transmembrane transport, catalytic activity, and membrane-associated cellular components (Figure 3C). The candidate strain-specific genes of CGRA01V4 include transporters that may be involved in nutrient acquisition and detoxification, ankyrin repeat-containing proteins, and fungal-specific transcription factors, suggesting a capacity for adaptation and gene regulation. Although the functions of the candidate strain-specific genes in CRO-I-41 are less diverse, they include dehydrogenases and binding proteins. ARG-5133-7 has a narrower range of specific functions but contains genes related to binding proteins and peptidases. Strain BR-85925-1 has genes in categories such as dehydrogenases, hydrolases and transporters, which may be associated with metabolic diversification. Finally, strain F-64330-2 shows a high presence of genes in functions such as dehydrogenases and hydrolases, suggesting potential roles in compound degradation. Other represented functional categories included fungal-specific transcription factors, oxidases and kinases, indicating differences in gene regulation and cell signaling functions among strains (Supplementary Table S8, Supplementary Figure S3).

### 4.5. Candidate strain-specific genes show similarity to known pathogenicity-related genes

Among the 733 candidate strain-specific genes of CGRA01V4, some had predicted functions potentially related to pathogenicity. However, 77% lacked functional annotation. Of the subset of candidate strain-specific genes affected by predicted high-impact effects, 92% had no known function and the remaining genes were classified as hydrolase, acetyltransferase, oxidase, and zinc finger. High-impact variants were detected in 50–90% of the candidate strain-specific genes assigned to the facilitator transporter, dioxygenase, WD-repeat, and oxidoreductase categories. High-impact variants were also detected in 39% of ankyrin-repeat-containing genes and 33% of ABC transporter genes, whereas 20–30% of genes assigned to the methyltransferase, fungal rhodopsin, fungal fatty acid synthase, tetratricopeptide-repeat, and cytochrome P450 categories carried predicted high-impact variants (Supplementary Table S8).

All *C. graminicola* strains evaluated showed necrotic lesions on maize, confirming their pathogenicity under the assay conditions (Figure 4C, D). However, lesion size differed significantly among strains (*F(4, 54.83)* = 12.71, *P* = 2.20 × 10^-7^). CRO-I-41 presented greater virulence than CGRA01V4 (Figure 4D), consistent with previous observations (Rogério et al., 2023). Based on this phenotypic contrast, the candidate strain-specific genes of these two strains were compared by performing blast searches against PHI-base. To focus specifically on plant pathogenicity-related associations, the PHI-base results were filtered to include only plant-associated pathogens. Of the 733 candidate strain-specific genes in CGRA01V4, 93 showed similarity to genes associated with plant pathogenicity phenotypes. Among these, 33 corresponded to genes for which gene disruption or deletion has been reported to cause reduced virulence or loss of pathogenicity. For CRO-I-41, 27 out of 200 candidate strain-specific genes were associated with plant pathogenicity-related entries in PHI-base, including 8 linked to reduced virulence and loss of pathogenicity phenotypes following gene disruption in previous studies. These genes represent candidates for future functional investigation of virulence in CRO-I-41.

### 4.6. Association patterns of INDELs, DUPs and LTR retrotransposons with plant-pathogen interaction genes in *C. graminicola*

The final dataset integrates 196,906 SVs and 13,565 TE annotations, encoded at the gene level as binary (presence/absence), count, and proportion-based variables within fixed 10 kb upstream and downstream flanking regions. Genome-wide sequence composition was assessed using 10 kb regions flanking genes across the genome (11,491 regions).

Genome-wide relationships among SVs, TEs, and genomic regions were examined to assess their associations with gene classes involved in plant interactions (Figure 5; Supplementary Table S4) revealing a non-uniform distribution of structural variants across genomic regions (Figure 5A). INDELs were significantly enriched in the 10-kb flanking regions upstream and downstream of plant-pathogen interaction genes, whereas their frequency within genes was comparatively lower (*Z=13.16; p=0.0001*). This pattern indicates that small-scale structural variation was more frequent in the flanking regions than within coding sequences. DUP and HDR (highly diverged regions) events also exhibited increased presence in flanking regions, although their distributions were less distinct than those of INDELs. The comparison between gene functional categories (see Figure 5B) showed that secreted and secreted-PHI-base genes showed a higher percentage of INDELs in upstream and downstream regions than non-secreted genes.

**Figure 5.**
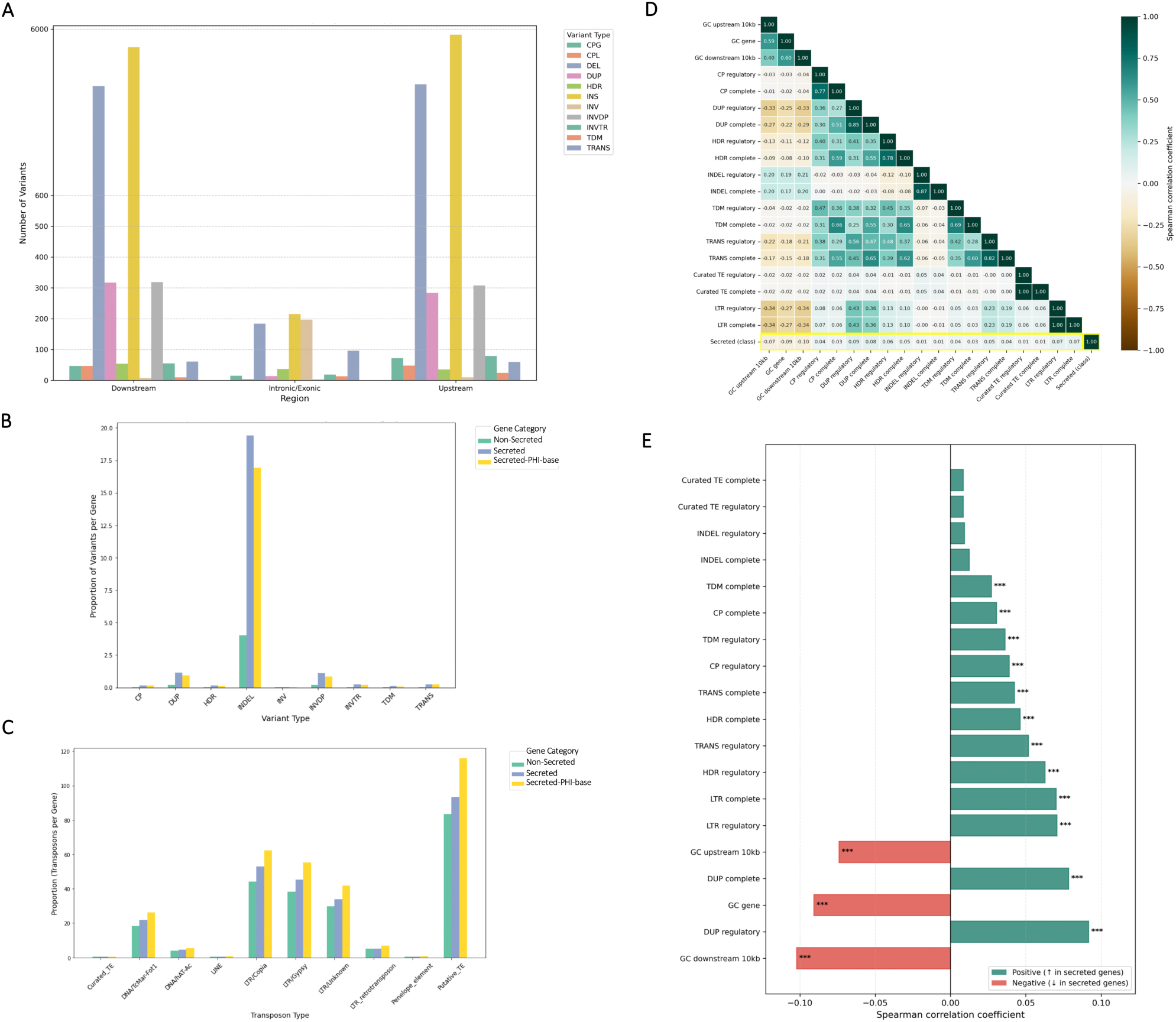
Genomic distribution of structural variants (SVs), transposable elements (TEs) and their association with plant-pathogen interaction genes. A) SV types across genomic regions (upstream, gene regions, and downstream). Bars represent the number of variants of each SV type, revealing a non-uniform distribution with a marked enrichment of INDELs, DUPs and HDR (highly diverged regions) events in potentially regulatory flanking regions. B) Proportion of structural variants normalized by gene category (Non-Secreted, Secreted, Secreted-PHI-base). Secreted and Secreted PHI-base genes exhibited a higher frequency of INDELs compared with non-secreted genes. C) Normalized counts of TEs per gene, grouped by TE superfamily and gene category (Non-secreted, Secreted and Secreted PHI-base). Secreted and Secreted PHI-base annotated genes show higher TE densities, particularly for LTR retrotransposons and DNA transposons. D-E) Spearman correlation coefficients describing the association between genomic features and the secreted gene class. D: Rank-based correlation heatmap depicting relationships among GC content, SV proportions in potentially regulatory flanking regions (10 kb upstream and downstream) and complete regions (regulatory and gene), as well as TE features. Correlations with the secreted gene class (positive class) are highlighted, providing a genome-wide overview of feature associations. E: Ranked Spearman correlation coefficients for the secreted gene class, extracted from panel D, highlighting the top positive and negative genomic features associated with secreted genes. Bar colors indicate the direction of the association, and statistical significance is denoted by asterisks.

TE density per gene also differed among gene categories, with secreted and secreted-PHI-base genes showing higher normalized counts of LTR retrotransposons and DNA transposons than non-secreted genes. This enrichment was stronger for LTR elements, indicating that LTR-containing regions are preferentially associated with secreted and Secreted-PHI-base genes (Figure 5C). The Spearman correlation analysis supported these observations, showing positive correlations with the secreted gene class for *DUP_complet*e, *DUP_regulatory*, *HDR_regulatory*, and *LTR_complete* features (Figure 5D), with DUP-related variables exhibiting the strongest positive associations. This pattern suggests coordinated enrichment of large-scale SVs, particularly DUP events, together with the presence of LTR retrotransposons. In contrast, GC content in both the upstream and downstream 10 kb regions showed significant negative correlations with secreted genes (Figure 5E), indicating that these genes tend to localize to relatively AT-rich genomic regions. The consistent directionality across regulatory and complete region features supports the presence of a compositional and structural bias associated with the secreted genes.

To further evaluate the relative contributions and interactions of these genomic features, a machine learning framework based on XGBoost modeling and SHAP analysis was applied (Figure 6). Large-scale INVs were excluded from this analysis due to the increased false-positive signals associated with chr04.

**Figure 6.**
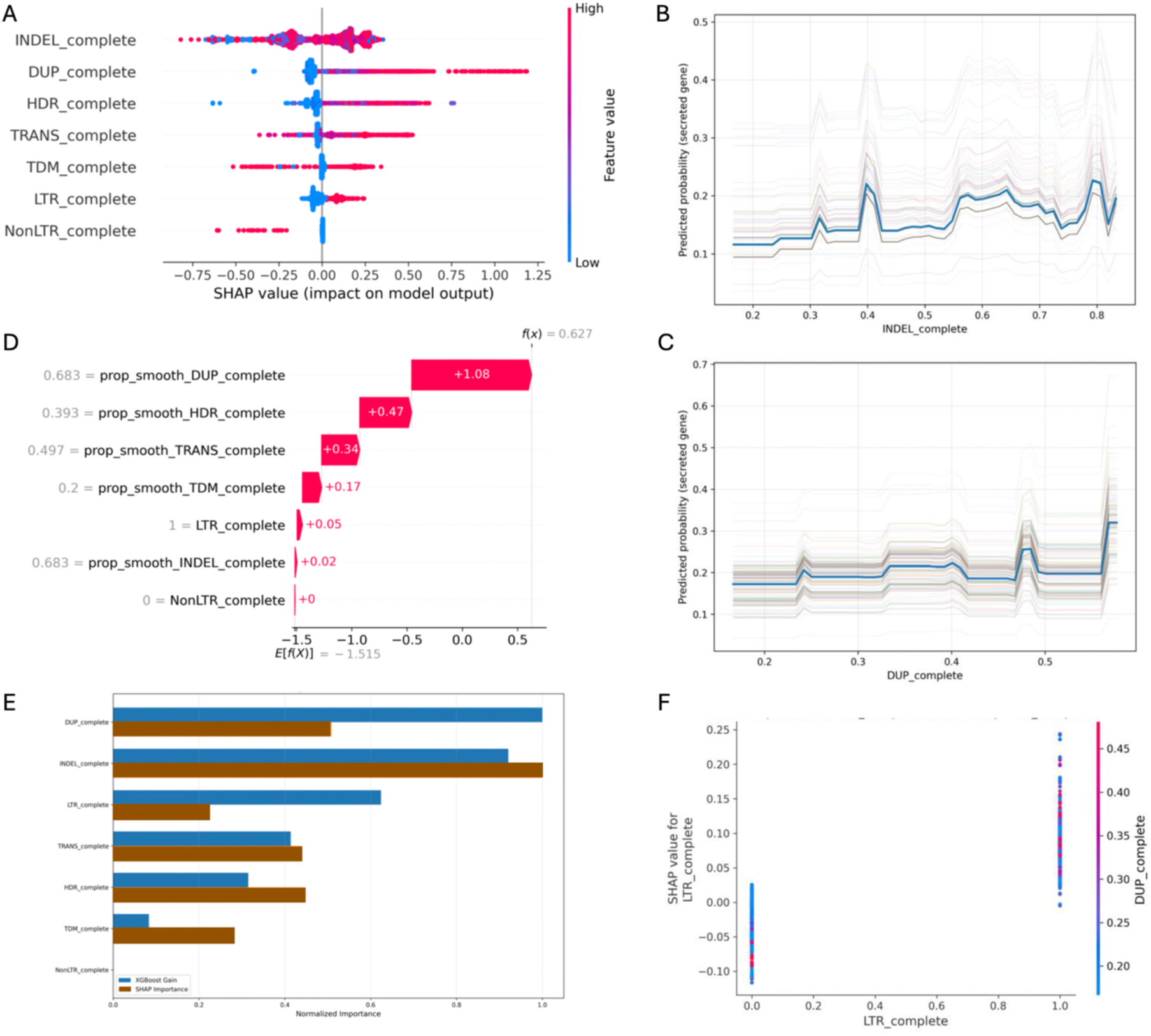
Genomic feature importance associated with secreted genes revealed by interpretable machine learning. A) SHAP summary plot showing the contribution and direction of genomic features associated with the secreted gene class. Each point represents a gene. The *y-axis* lists genomic features, while the *x-axis* shows SHAP values, indicating whether each feature contributes positively or negatively to the association with secreted genes. Colors represent feature values, ranging from low (blue) to high (pink), illustrating how feature levels influence the direction and magnitude of their association with the secreted gene class. B-C) Partial dependence and individual conditional expectation (PDP–ICE) plots illustrating the relationships between complete-region structural-variant occurrence proportions and the model-predicted probability of the secreted-gene class. Panel B shows *INDEL_complete*, where increasing *INDEL_complete* values are associated with changes in the model-predicted probability of the secreted-gene class. Panel C shows *DUP_complete*, revealing a stepwise and non-linear relationship between *DUP_complete* values and the model-predicted probability of the secreted-gene class. D) SHAP waterfall plot showing the local feature contributions for a selected true-positive secreted gene. Values are shown on the XGBoost raw-margin scale: the expected value (−1.515) plus the displayed feature contributions equals the model output (0.627; predicted probability = 0.652). Positive contributions move the model output toward the secreted-gene class, with *prop_smooth_DUP_complete* providing the largest positive contribution in this example. E) Comparison of feature importance rankings obtained from XGBoost gain and mean absolute SHAP values. Both approaches consistently identify DUP-, INDEL-, and LTR-related features as the most influential genomic variables associated with secreted genes. F) SHAP dependence plot for *LTR_complete*, colored by *prop_smooth_DUP_complete*, illustrating the relationship between LTR retrotransposon presence and duplication context. *LTR_complete* is a binary variable (presence/absence), whereas *prop_smooth_DUP_complete* represents the smoothed proportion of available strain comparisons in which a duplication was observed within the complete gene-associated region. Each point represents a gene, and color intensity reflects increasing duplication levels. The plot shows that, for genes with LTR presence (LTR = 1), higher duplication values (represented by warmer colors) are associated with a stronger contribution toward the secreted gene class, indicating modulation by the local duplication context.

Table 2 summarizes the performance of the XGBoost models across the three evaluated feature blocks. The first metric shows the ROC-AUC obtained on the held-out test set. The second reports the mean ROC-AUC from five-fold cross-validation grouped by contig and performed on the training set, together with its 95% interval derived from variability across folds. The remaining columns report, in this order, F1, precision, recall, and average precision (AP), all calculated on the held-out test set. ROC-AUC on the held-out test set ranged from 0.600 to 0.646. In cross-validation grouped by contig, mean ROC-AUC ranged from 0.598 to 0.635, and the lower bounds of the fold-based 95% intervals remained above 0.5 in all three configurations. F1 ranged from 0.208 to 0.257; precision from 0.122 to 0.176; recall from 0.454 to 0.699; and AP from 0.148 to 0.214. The reduced Feature Block 3 achieved the highest recall (0.699) but the lowest precision (0.122) and ROC-AUC (0.600), indicating that the reduced feature set increased the identification of positive cases (secreted genes) at the expense of precision and overall discriminatory performance relative to Blocks 1 and 2. To assess whether this signal and the identified associations were maintained across changes in the contig partition, the analysis was repeated using 30 different `GroupShuffleSplit` seeds. Across all three blocks, the lower bound of the central 95% percentile interval of the ROC-AUC distributions remained above 0.5. SHAP feature-importance rankings also showed high concordance across repetitions (Kendall’s W = 0.91– 0.94). These results show that the discriminatory signal was reproducible and that the feature hierarchy was not driven by a particular partition of the data.

**Table 2.** Performance of the XGBoost models across the three feature-block configurations.

| Variable | Feature Block 1 | Feature Block 2 | Feature Block 3 |
| --- | --- | --- | --- |
| Test ROC-AUC | 0.646 | 0.634 | 0.600 |
| CV ROC-AUC, mean [95% CI] | 0.635 [0.617–0.653] | 0.614 [0.584–0.643] | 0.598 [0.572–0.623] |
| F1 | 0.257 | 0.236 | 0.208 |
| Precision | 0.176 | 0.159 | 0.122 |
| Recall | 0.474 | 0.454 | 0.699 |
| AP | 0.214 | 0.174 | 0.148 |

CV, cross-validation; CI, confidence interval; AP, average precision.

The SHAP summary plot ranked *DUP_complete* and *INDEL_complete* among the strongest contributors to the secreted-gene class (Figure 6A). PDP and SHAP dependence analyses revealed non-linear response patterns for both variables. SHAP values for *INDEL_complete* showed that its high overall importance did not result from a steady increase in model contribution as *INDEL_complete* values increased (Figure 6B). Similarly, *DUP_complete* showed a stepwise relationship with the model contribution to the secreted-gene class (Figure 6C). These patterns indicate that the associations captured by the model vary across specific ranges of these genomic variables and may also depend on other genomic features. Notably, *INDEL_complete* ranked first by SHAP importance in Feature Block 3 across all 30 resamples, but it was not significantly associated with the secreted-gene class in the univariate analysis (*Spearman’s rho* = 0.012, *P* = 0.12). Its SHAP contribution varied across *INDEL_complete* values and differed depending on DUP. Therefore, its high importance within the model should not be interpreted as evidence that increasing INDEL proportions consistently increase or decrease the probability of a gene belonging to the secreted-gene class. For the selected true-positive gene, the SHAP waterfall plot showed that the DUP feature made the largest positive contribution to the model output associated with the secreted-gene class (Figure 6D). Similar rankings obtained using XGBoost gain and SHAP importance further supported the importance of these variables within the model (Figure 6E; Supplementary Figure S4). Notably, the SHAP dependence analysis showed that the model-learned contribution associated with LTR presence varied according to local DUP proportions (Figure 6F). In regions containing LTR retrotransposons, higher DUP proportions were associated with stronger positive SHAP contributions to the secreted-gene class, whereas this pattern was weaker in regions without LTRs. These results indicate that the model-learned association between DUP features and the secreted-gene class varied according to the presence of LTR retrotransposons. However, this model-based association does not demonstrate that LTR retrotransposons caused the duplication events.

## 5. Discussion

We present genome sequences of four *C. graminicola* strains, representing three geographically isolated genetic lineages, sequenced using both short and long reads technologies. We obtained chromosome-level assemblies for strains CRO-I-41, ARG-5133-7, and near chromosome-level assemblies for BR-85925-1 and F-64330-2. Comparison of these genome sequences with the reference genome (CGRA01V4) revealed eight highly syntenic core chromosomes, two core chromosomes exhibiting substantial large-scale SVs, including inversions and translocations, and variable minichromosomes that differ in number among strains. All four strains encoded similar numbers of CAZymes, secreted proteins, and potential effectors, comparable to those identified in the reference genome. These protein groups contribute to diverse enzymatic activities and pathogenic mechanisms in fungal plant pathogens (Baroncelli et al., 2024; Kleemann et al., 2012; Stergiopoulos and De Wit, 2009). The relatively limited variation observed among strains is consistent with the findings of Gan et al. (2016), who reported broadly similar CAZyme repertoires among *Colletotrichum* species, while also suggesting that differences in CAZyme profiles may reflect adaptation to specific infection.

The synteny analysis revealed the genomic organization of *C. graminicola*, indicating a clear differentiation between core and minichromosomes. We found higher levels of synteny across strains for the core chromosomes, indicating a conserved genetic core. Similar patterns have been reported in other plant-pathogenic fungi in which core chromosomes generally show high levels of synteny across strains (Badet et al., 2021; Gan et al., 2021; Lapalu et al., 2024; Zaccaron et al., 2022). Chr02 showed more than 93% synteny (Figure 1A). In contrast, chr04 and chr09 showed lower levels of synteny than the other core chromosomes, while the minichromosomes exhibited markedly reduced synteny (32.48%) (Figure 1C).

Our results revealed various types of genomic variation, including translocations, duplications, and inversions distributed throughout the genome. The distribution of structural variants is consistent with differential evolutionary constraints across genomic regions, with gene regions being less affected, suggesting stronger conservation of genes. Interestingly, most structural variants were located in intergenic regions, suggesting that these changes may not directly alter the coding sequences but instead could influence gene regulation. Rech et al. (2014) found that intergenic regions in *C. graminicola*, including non-coding sequences, are under positive selection. They found that several of these regions were associated with gene clusters involved in pathogenicity. Specifically, they identified a pattern in which 5′ UTR regions regulate pathogenicity-related genes, thereby influencing phenotypic plasticity. The authors also suggested that structural variation among isolates results from positive selection. Overall, these findings indicate that these intergenic regions are likely shaped by selective pressure acting on the genome.

Segmental duplications in CGRA01V4, particularly involving the core chr04 and minichromosomes 11 and 13, may represent an additional source of genome structural variability in *C. graminicola*, similar to that reported in *Magnaporthe oryzae* (Langner et al., 2021). Although we did not identify genes in the strain-specific regions containing segmental duplications, as has been found in other *Colletotrichum* species (Lapalu et al., 2024), we confirmed the occurrence of segmental duplications between a core chromosome and two minichromosomes. This pattern of segmental duplication, while lacking gene content, could still play an important role in genome evolution. Such duplicated regions could provide genomic substrate for evolutionary innovation (Andersson et al., 2015; Fouché et al., 2018). In addition, genomic rearrangements, such as translocations and inversions directly alter chromosome structure and organization and can affect the expression of nearby genes, including effector genes and other virulence-associated genes, in pathogenic fungi. Repetitive elements and TEs can also promote such rearrangements and affect gene expression, as reported in *Verticillium dahliae* (Faino et al., 2016) and *Zymoseptoria* species (Lorrain et al., 2021).

Our comparative analysis shows clear structural variability in the minichromosomes among *C. graminicola* strains. In CGRA01V4, these chromosomes also show a higher variant density than the core chromosomes, together with low synteny, high repeat content (Becerra et al., 2023), and a strong enrichment in candidate strain-specific genes. In addition, minichromosomes 11 and 13 are involved in segmental duplications with core chr04. Similar characteristics have been reported for minichromosomes in other *Colletotrichum* species, where they are often repeat-rich and gene-poor (Becerra et al., 2023; Gan et al., 2021; Pires et al., 2016; Wang et al., 2023). However, their gene content and their possible contribution to pathogenicity differ among species. In several *Colletotrichum* species, minichromosomes and other accessory chromosomes carry host-specific effector or virulence genes, as in the dispensable chromosome required for virulence in *C. higginsianum* (Plaumann et al., 2018), the virulence-governing minichromosome of *C. lentis* (Bhadauria et al., 2019), and the pathogenesis-related minichromosome genes of mango-infecting *C. asianum*, whose disruption reduces virulence (Wang et al., 2023); predicted effectors, CAZymes and biosynthetic gene clusters have similarly been reported on the large accessory chromosomes of *C. cereale* (Cooper et al., 2026). In contrast, the minichromosomes of the *C. graminicola* reference strain are gene-poor and contain no predicted secreted protein-encoding genes or candidate effectors, suggesting that their possible contribution to genome variability is not associated with an effector-rich minichromosome compartment. A directly comparable situation occurs in *C. lupini*, whose dispensable minichromosome (chromosome 11) lacks candidate effectors and is absent from some pathogenic isolates, indicating that it is not required for pathogenicity (Menicucci et al., 2026). In *C. graminicola*, the contribution of minichromosomes to genome variability may therefore be related to other features, particularly their repeat content, structural variation, and strain-specific gene content. In CGRA01V4, candidate strain-specific genes represent 28%, 93%, and 100% of the annotated genes on minichromosomes 12, 11, and 13, respectively. These differences suggest that minichromosomes and other accessory chromosomes do not play the same role in all *Colletotrichum* species. Rather, their contribution to pathogenicity or adaptation may depend on the genetic content of each lineage, supporting an emerging picture in which these chromosomes can contribute to genome evolution through different mechanisms and are not necessarily dedicated effector reservoirs.

Candidate strain-specific genes were located within highly dynamic genomic regions, which has also been shown by Croll and McDonald (2012) and De Jonge et al. (2013) in other species of plant pathogenic fungi. The variable range of candidate strain-specific genes found in this study has also been reported in other fungi, including *Colletotrichum* spp., where between 31 and 731 such genes have been described (Wang et al., 2023). When we considered their evolutionary relationships (Rogério et al., 2023), strains belonging to the European lineage had similar numbers of candidate strain-specific genes, whereas the Brazilian and North American representatives showed higher numbers. These values are consistent with the genetic diversity and genomic history of these lineages. For instance, the North American lineage (here represented by CGRA01V4) has a higher genetic diversity and appears to have served as an intermediate between an unknown source population and the rest of the world (Rogério et al., 2023). Thus, the differences in the number of candidate strain-specific genes may reflect the evolutionary divergence among strains. CRO-I-41 and ARG-5133-7, which showed the highest numbers of variant observations relative to CGRA01V4, also differed significantly in virulence in the infection assays. This association may indicate that genomic variation is related to differences in virulence among strains.

Interestingly, many candidate strain-specific genes identified in CGRA01V4 have homologous genomic sequences in other strains but no corresponding gene model. Some of these loci also contain high-impact variants predicted to disrupt gene function. These observations suggest that some loci classified as strain-specific may represent lineage-specific pseudogenization rather than true gene absence or recent gene acquisition. Pseudogenes provide evidence of gene loss during genome evolution (Lafontaine and Dujon, 2010; Van Der Burgt et al., 2014). Similarly, lineage-specific pseudogenization has been reported in *Fusarium graminearum*, highlighting gene decay as an important component of pathogenic genome evolution (Rau et al., 2025). Pseudogenes may also contribute to genome evolution through processes such as gene conversion and recombination with functional homologs (Balakirev and Ayala, 2003; Prade et al., 2018; Yadav et al., 2023). However, the biological activity of these disrupted loci cannot be inferred from sequence data alone. Future transcriptomic and functional analyses will be needed to determine their potential roles and clarify the contribution of lineage-specific pseudogenization to the diversification of strain-specific gene repertoires.

Comparison of the candidate strain-specific genes from CGRA01V4 against PHI-base identified homologs with experimentally demonstrated roles in host-pathogen interactions in other fungi. Comparative analysis of these strain-specific genes with homologous genes characterized in other plant pathogens, such as *C. higginsianum* (O’Connell et al., 2012) and *Magnaporthe oryzae* (Dean et al., 2005) has provided functional context for these candidate genes. A detailed functional gene description and supporting literature are provided in Supplementary Note 1. These candidate strain-specific genes include functions related to metabolism, transport, transcription regulation, and effector proteins. Previous transcriptional profiling showed some are expressed during the early stages of biotrophy (O’Connell et al., 2012). Several candidate strain-specific genes expressed during early biotrophy showed similarity to genes whose disruption in other fungi affects appressorium formation or infection establishment. Comparative transcriptomic and functional analyses could help identify candidate genes involved in plant colonization by *C. graminicola*.

To better understand the genomic context of secreted genes in *C. graminicola*, we used an integrative machine learning approach to evaluate associations between genomic features and secreted genes. Rather than serving as the primary evidence or as a predictive or causal framework, the XGBoost analysis provided multivariate support for the descriptive and correlation-based patterns presented above. SHAP consistently identified INDELs, duplications, and LTR retrotransposons among the most informative features associated with secreted genes across chromosome-grouped partitions. Machine learning approaches have also been applied to fungal genomic data, although with different analytical objectives. At a broad evolutionary scale, Fijarczyk et al. (2025) identified gene content, including effector repertoires, as an important genomic correlate of pathogenic lifestyle across Sordariomycetes, whereas at a finer genomic scale, gradient-boosting models have been used to examine associations between local genomic features and the occurrence of SVs in *Zymoseptoria tritici* (Badet et al., 2021). The consistency of SHAP feature rankings across chromosome-grouped partitions supported the stability of the relative importance assigned to these features. Although the overall discriminatory performance of the models was moderate, our analysis was not intended solely as a predictive framework. By combining XGBoost with SHAP, we identified consistent, non-linear, and range-dependent associations between genomic features and secreted genes that would not be adequately captured by simple univariate analyses. To our knowledge, this is a novel application of interpretable machine learning to investigate how structural and repetitive genomic features associate with secreted genes in a fungal plant pathogen.

INDELs were enriched in the upstream and downstream flanking regions of genes encoding secreted proteins, which may have potential regulatory relevance. The importance assigned by the XGBoost model to *prop_smooth_INDEL_complete* represents a related but distinct result, because this variable captures the proportion of the complete gene-associated region affected by INDELs, which the model evaluates together with other genomic features. Therefore, the enrichment of INDELs in these flanking regions does not imply that increasing INDEL density progressively increases the probability of a gene belonging to the secreted-gene class. Rather, the model indicates a more complex, non-linear relationship that also depends on other genomic features.

Similar associations between structural variation and genes involved in pathogenicity or adaptation have been reported in other fungal plant pathogens. In *Fusarium graminearum*, deletions, translocations, and other structural variants affect genomic regions associated with secondary metabolism and effectors (Dhakal et al., 2024; Guo et al., 2021). In *Verticillium dahliae*, repeat-associated rearrangements, gene duplications, differential gene loss, and polymorphic transposable elements contribute to dynamic genomic regions containing virulence-related genes and have been associated with variation in the expression of nearby pathogenicity-related genes (Faino et al., 2016; Torres et al., 2021). Likewise, in *Rhynchosporium commune*, copy-number variation preferentially affects genes involved in host exploitation, including predicted effectors and cell wall-degrading enzymes (Stalder et al., 2023). Together, these studies support a broader association between structural genome variation and the diversification of genomic regions involved in fungal pathogenicity and adaptation.

Interestingly, our analysis identified a co-occurrence between LTR retrotransposons and duplication-related features in genomic regions associated with secreted genes, supporting a shared structural context that may contribute to genome plasticity. Similar associations between TEs, genomic rearrangements, and duplicated regions have been reported in *V. dahliae* (Faino et al., 2016) and *Fusarium oxysporum* (Hua-Van et al., 2000). In *Colletotrichum* spp., *Gypsy*-type LTR retrotransposons have also been associated with secretome composition, effector genes, and secondary metabolism gene clusters (Gan et al., 2021). TEs can promote genome reorganization through mechanisms such as double-strand breaks and subsequent repair, although their effects vary among fungal lineages depending on TE composition and genomic context (Sauters and Rokas, 2025). Within this broader context, our results identify a more specific pattern in *C. graminicola*, in which LTR retrotransposons and duplications tend to co-occur in genomic regions associated with genes involved in plant-pathogen interactions. This co-occurrence is consistent with dynamic genome organization, although the mechanisms underlying this association remain unknown.

These results show that structural variants, including INDELs within potentially regulatory flanking regions and duplications, as well as transposable elements such as LTR retrotransposons, are associated with the distribution and genomic context of plant-pathogen interaction genes in *C. graminicola*. This characterization underscores the complex genomic landscape associated with pathogen virulence-related traits, offering new hypotheses for future functional validation studies. In addition, this approach opens the possibility of incorporating a broader set of variables and data layers, including transcriptomic, functional, and population-level information, which when analyzed together, may help to understand the complexity of genome organization and structural variation associated with pathogenicity.

Overall, these findings expand our knowledge of the *C. graminicola-*maize pathosystem by linking structural variation, transposable elements, and genome organization with plant-pathogen interaction genes. They also provide testable hypotheses for future functional studies of anthracnose pathogenicity and fungal genome evolution. In practical terms, the prevalence of patterns consistent with lineage-specific pseudogenization implies that effector and virulence-gene catalogues derived from a single reference genome may misrepresent the functional repertoire of field strains, while the strain-specific candidate virulence factors identified here, for example in the highly virulent strain CRO-I-41, provide concrete targets for functional validation and for more durable anthracnose management.

## 7. Author Contributions

SB, MRT, and SAS conceived and designed the study. SB, PGR, LRM and SGS performed laboratory experiments and generated biological material. SB and JM performed the machine learning analyses. SB, TRB, JLVV and LF performed the statistical and bioinformatic analyses. SB wrote the manuscript. PGR, JM, SAS, and MRT contributed to review and editing of the manuscript. PGR, JM, SGS, FBCF, FR, TRB, LF, RB, SAS, and MRT contributed to data interpretation and provided critical revisions. All authors read, revised, and approved the final version of the manuscript.

## 9. Acknowledgments

The authors thank the Supercomputing and Bioinnovation Center (SCBI) of the University of Málaga for their provision of computational resources and technical support (http://www.scbi.uma.es/site). The Galaxy server that was used for some calculations is in part funded by Collaborative Research Centre 992 Medical Epigenetics (DFG grant SFB 992/1 2012) and German Federal Ministry of Education and Research (BMBF grants 031 A538A/A538C RBC, 031L0101B/031L0101C de.NBI-epi, 031L0106 de.STAIR (de. NBI)).

## 10. Data Availability Statement

The genome assemblies of the four strains used in this study can be found in the NCBI database under BioProject ID PRJNA1444001.

## 11. Extended data

**Supplementary Table S1.** Raw long-read sequencing statistics for each *Colletotrichum graminicola* strain.

**Supplementary Table S2.** Mapping of genomic features to the feature-block configurations used in the machine learning analyses

**Supplementary Table S3.** Biological classification and metadata of genomic features used in the machine learning analyses.

**Supplementary Table S4.** Master gene-level dataset integrating structural variants, transposable elements, sequence composition, and gene-class information used for machine learning analyses.

**Supplementary Table S5.** Characteristics of minichromosomes in strains with chromosome-level genome assemblies.

**Supplementary Table S6.** Structural variants identified in the CGRA01V4 reference genome in pairwise comparisons with the other *Colletotrichum graminicola* strains.

**Supplementary Table S7.** Functional annotations of CGRA01V4 genes affected by at least one predicted high-impact effect in the four pairwise strain comparisons.

**Supplementary Table S8.** Functional annotations of candidate strain-specific genes in five *Colletotrichum graminicola* strains.

