## Supplementary Figures for "Genome-wide structural variants uncover strain-specific genes and evolutionary genomic determinants of a fungal maize pathogen"

**Supplementary Figure S1.** plotsr visualization of SyRI-defined syntenic regions and large-scale structural rearrangements between the reference genome CGRA01V4 and each of the four *Colletotrichum graminicola* strains. In CGRA01V4, contigs correspond to chromosomes.

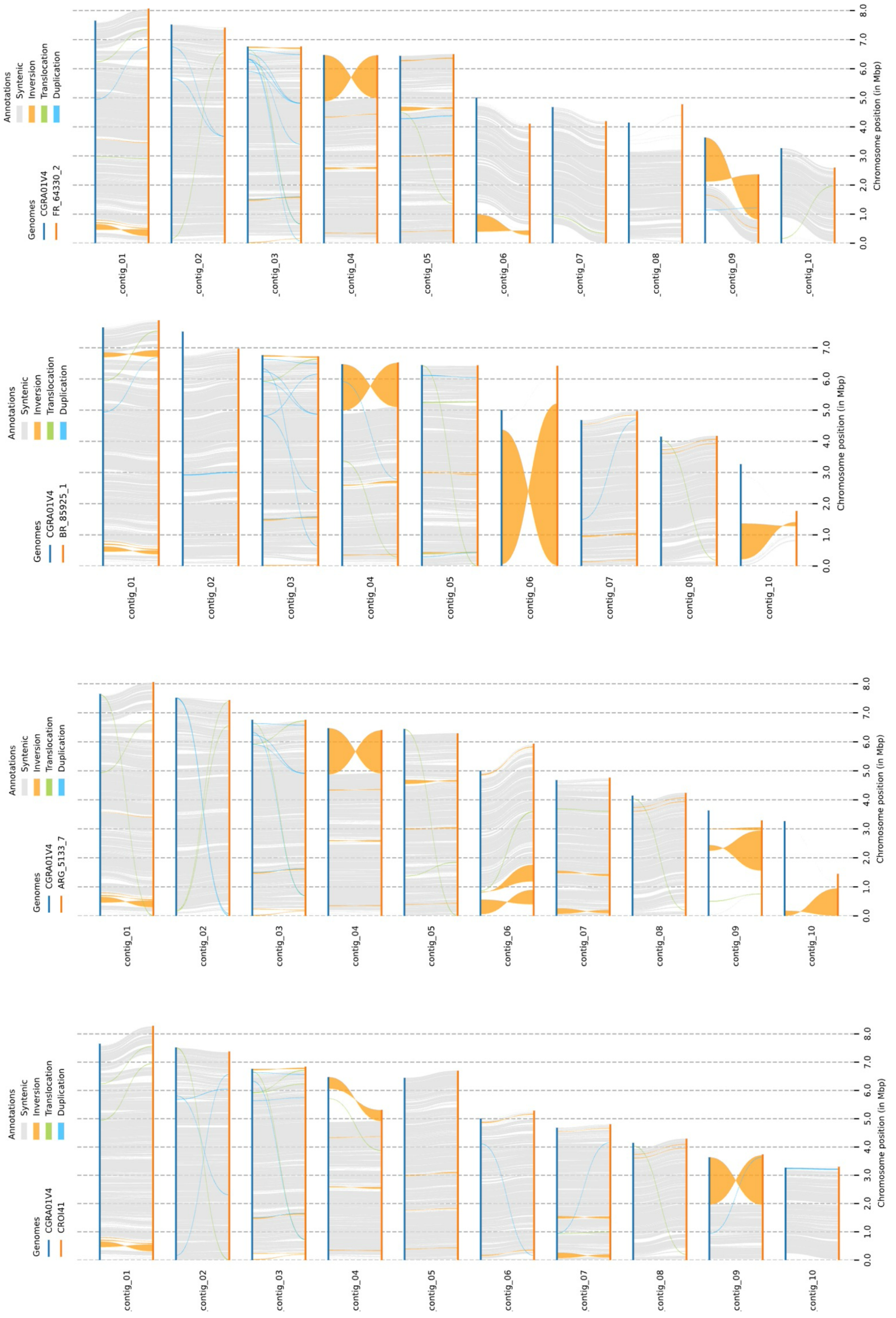

**Supplementary Figure S2.** Circular genome plots showing segmental duplications in the reference genome CGRA01V4 and the four *Colletotrichum graminicola* strains CRO-I-41, ARG-5133-7, F-64330-2, and BR-85925-1. Segmental duplications were detected in all strains; however, the specific duplicated regions (~174 kb) linking core chr04 with minichromosomes chr11 and chr13 were unique to CGRA01V4. Detailed characterization of these strain-specific duplicated regions was therefore focused on CGRA01V4.

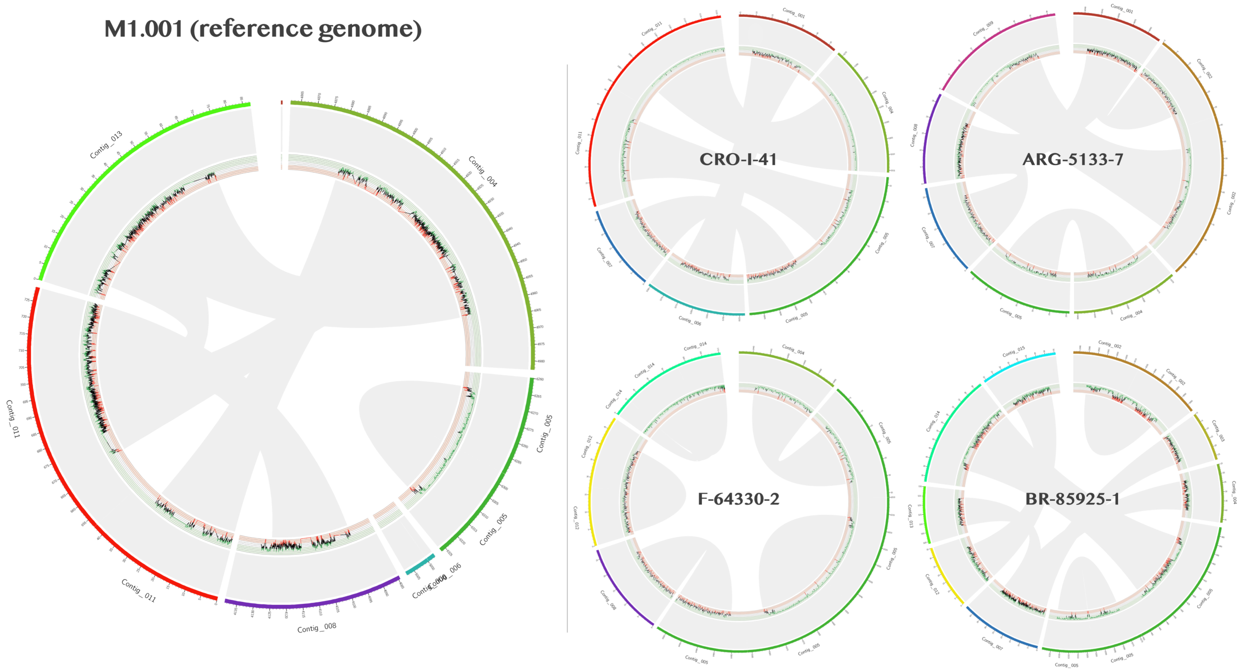

**Supplementary Figure S3.** Distribution of functional categories among candidate strain-specific genes in *Colletotrichum graminicola* strains. Bars show the number of candidate strain-specific genes assigned to each functional category in each strain.

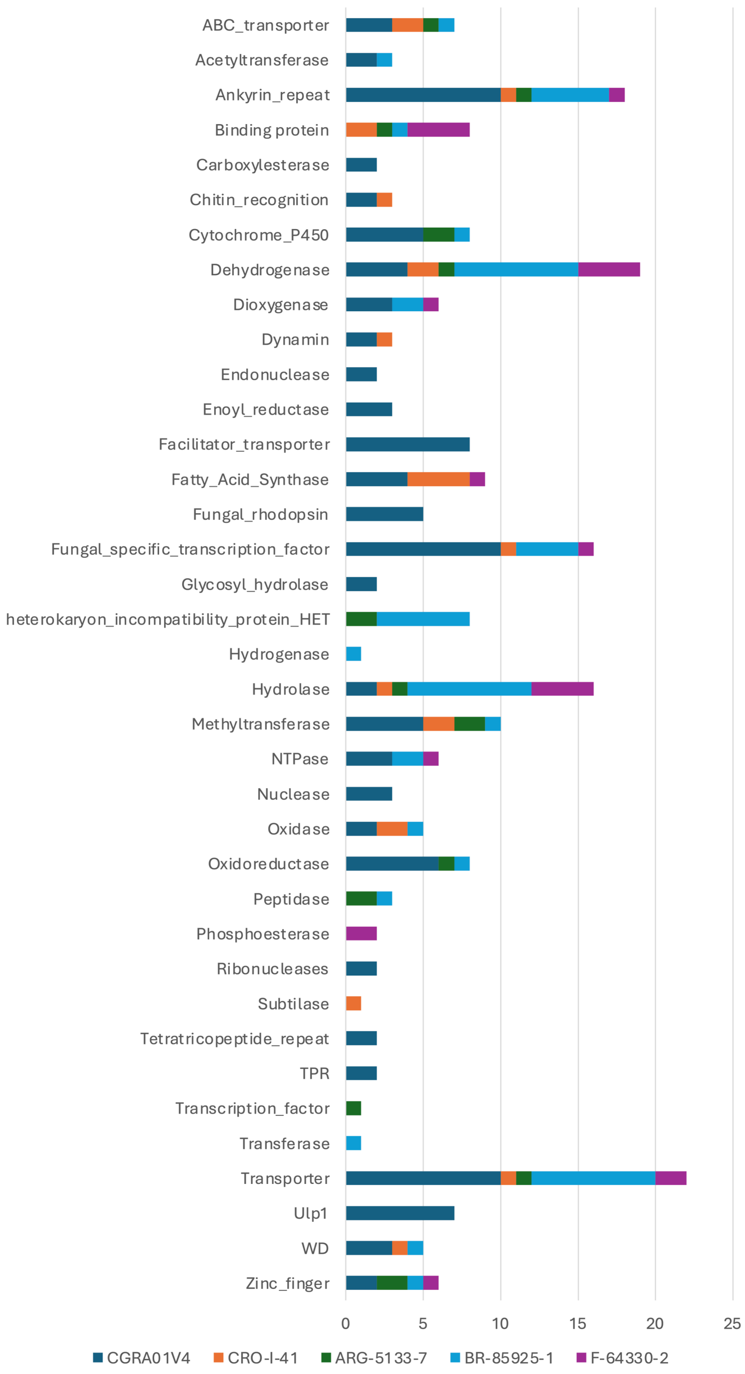

**Supplementary Figure S4.** Comparison of feature importance across the three feature blocks. Normalized feature importance derived from XGBoost gain (blue) and mean absolute SHAP values (orange) is shown for each feature block.

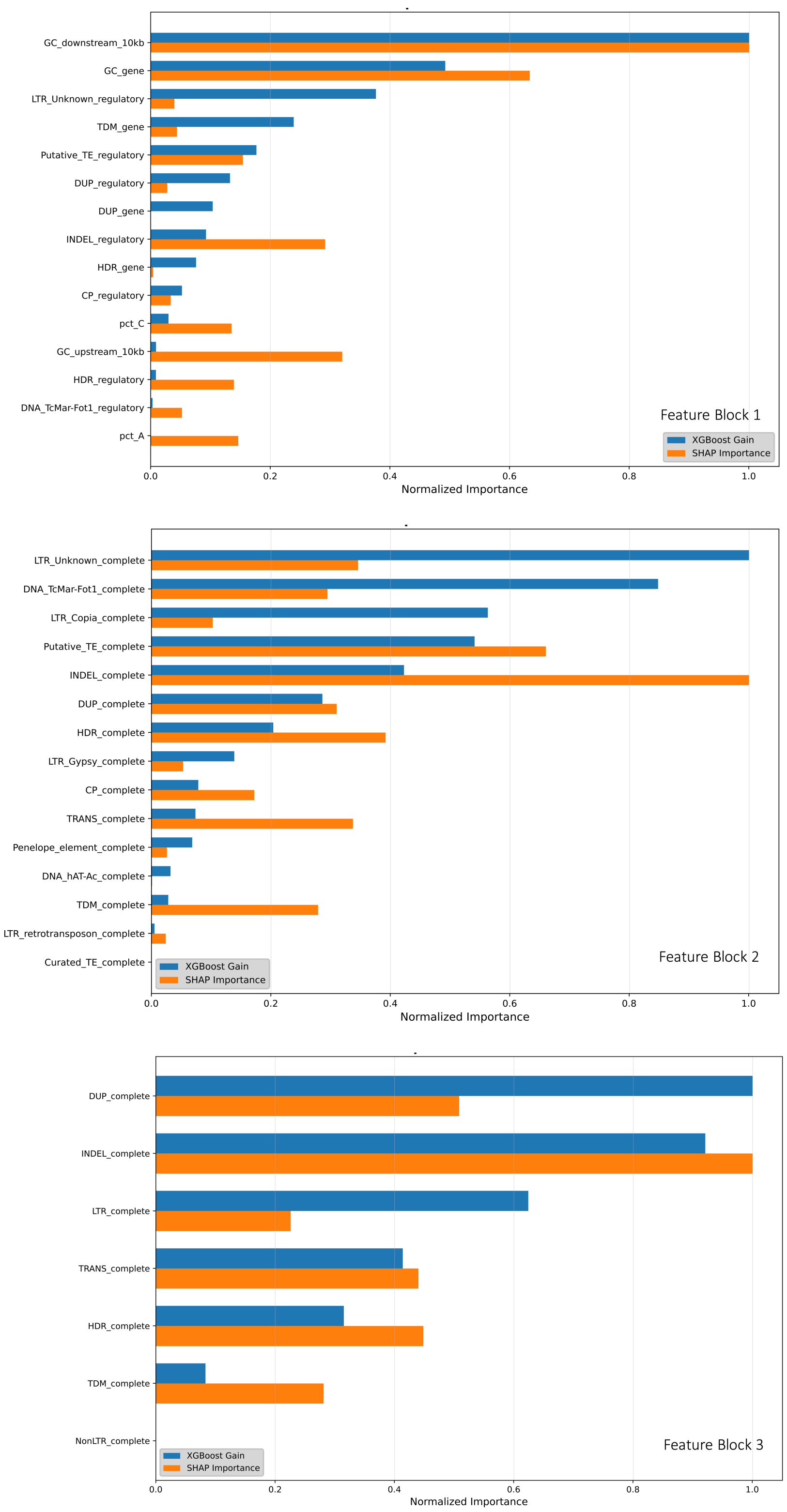
