## Supplementary Note S1 for "Genome-wide structural variants uncover strain-specific genes and evolutionary genomic determinants of a fungal maize pathogen"

### 1. Functional description of candidate strain-specific genes based on PHI-base and homology analysis

Comparison of the candidate strain-specific genes from CGRA01V4 against PHI-base identified homologs with experimentally demonstrated roles in host-pathogen interactions in other fungi. Comparative analysis of these strain-specific genes with homologous genes characterized in other plant pathogens, such as *C. higginsianum* (O'Connell et al., 2012) and *Magnaporthe oryzae* (Dean et al., 2005) has provided functional context for these candidate genes. Analysis of transporter homologs provides functional context for biological processes potentially associated with pathogenicity in *C. graminicola*. For example, CGRA01v4_12258 (MGG_05505) encodes the vacuolar flux protein ATG22, which is associated with autophagy, an important process for plant infection by *M. oryzae* during appressorium development (Osés-Ruiz, 2014; Osés-Ruiz et al., 2021; Veneault-Fourrey et al., 2006).

The ABC transporter *CGRA01v4_00253* (*GLRG_05467*), previously known as *CgrABCC1c*, shows differential expression patterns, with higher expression levels in appressoria than in the biotrophic phase, suggesting a specialized role in the pre-penetration phase of maize infection (Dube et al., 2018; O'Connell et al., 2012). Similarly, *CGRA01v4_13348* (*CH063_00247*), another ABC transporter identified as *ChiABCG4*, is highly expressed during the biotrophic stage of *Arabidopsis thaliana* infection, highlighting its importance in this phase (Dube et al., 2018; O'Connell et al., 2012). In addition, ABC transporter genes serve as virulence factors by assisting in detoxification (Oliveira et al., 2022).

Further investigation of sulfur metabolism identified *CGRA01v4_06040* (*MGG_03103*), which shared homology with the high-affinity methionine permease of

*Neurospora crassa*. Methionine permeases, although less characterized in fungi, are integral to sulfur metabolism (Paietta, 2014). In the context of carbon metabolism, several genes such as *CGRA01v4_07996* (*GLRG_02169*) and *CGRA01v4_13495* (*GLRG_11320*), annotated as alpha glucoside and hexose transporters, respectively, exhibit transposon insertions within these genes, potentially impacting their function (Braga et al., 2014). Additionally, the putative quinate permeases *CGRA01v4_14257* and *CGRA01v4_14259* (both *MGG_09778*) show differential expression during appressorium formation. These transporters allow quinate to enter the fungal cell of *M. oryzae*, where it can be metabolized, enabling its growth and development of the infection (Soanes et al., 2012). Another transporter, *CGRA01v4_14345* (*MGG_13562*), identified as a major facilitator superfamily sugar transporter, is expressed during appressorium maturation in *M. oryzae* infection. This pathogen may use sugar transporters within the appressorium to prepare for tissue invasion and to exploit host carbohydrates as a nutrient source (Soanes et al., 2012).

Among the enzymes and proteins identified, several belonged to the P450 family, including CGRA01v4_04683, CGRA01v4_01484, CGRA01v4_06137, and CGRA01v4_12023. This group of proteins belongs to the P450 monooxygenases complement, which, although not directly related to pathogenicity, may be involved in the biosynthesis of secondary metabolites and plant defense mechanisms (Ortiz-Álvarez et al., 2024). *CGRA01v4_06138* (*GLRG_11834*) is identified as CYP539C1 by Ortiz-Álvarez et al. (2024). Its homolog in *M. oryzae* (*MGG_12496*) is associated with xenobiotic metabolism and toxin biosynthesis, crucial during host colonization (Yan et al., 2023; Zhang et al., 2022). The role of detoxification is highlighted by peroxidases such as *CGRA01v4_01673* (*MGG_08046*) and *CGRA01v4_12556* (*MGG_07500*), which help pathogens detoxify host-derived reactive oxygen species (ROS) during infection

(Huang et al., 2017). Some genes related to lipid metabolism were found that could play a role during appressorium development, such as: *CGRA01v4_15038* (*MGG_06704*), which is related to phospholipid metabolism (Franck et al., 2015); *CGRA01v4_05870* (*MGG_10414*) triacylglycerol lipase VTL1, which takes part in fatty acids and glycerol metabolism (Hong et al., 2021; Wang et al., 2007).

Cell wall degradation is another critical process in fungal pathogenicity (Baroncelli et al., 2024). *CGRA01v4_14350* (*MGG_05125*) and *CGRA01v4_09717* (*MGG_01542*), encoding GH18 family CAZymes and GH10 family xylanases, respectively, play significant roles in degrading plant cell walls during the early stages of *M. oryzae*-rice infection (Du et al., 2024; Nguyen et al., 2011). Two chitin-related genes, *CGRA01v4_02842* (*GLRG_03690*), a chitinase, and *CGRA01v4_08062* (*GLRG_02208*), a chitin deacetylase, have CBM18 domains in their structure. In *M. oryzae*, CBP1 is a putative extracellular chitin-binding protein involved in surface sensing during appressorium differentiation (Kamakura et al., 2002). Secondary metabolism genes such as *CGRA01v4_01998* (*GLRG_08632*), may be related to melanin biosynthesis. *GLRG_08632*, a non-reducing polyketide synthase (PKS) is part of a cluster of genes producing unknown secondary metabolites that are expressed specifically during appressorium formation, prior to penetration (Torres, 2013; Torres et al., 2016). In addition, *CGRA01v4_06139* (*MGG_08236*) is annotated as a reducing PKS found in a cluster of secondary metabolite biosynthetic genes. This gene is exclusively active during pathogenesis and likely plays a crucial role in the colonization of the host (Mehta et al., 2023).

Transcription factors play an important role in regulating gene expression during infection (Torres et al., 2016). *CGRA01v4_13778* (*MGG_10212*), identified as *RPP1,* is part of a network of transcription factors regulating appressorium-mediated plant infection in *M. oryzae* (Osés-Ruiz et al., 2021). *CGRA01v4_01399* (*MGG_13350*), a Zn(2)-Cys(6) transcription factor, is upregulated during appressorium morphogenesis in *M. oryzae* (Osés-Ruiz, 2014; Osés-Ruiz et al., 2021). *CGRA01v4_13039* (*MGG_01836*), is classified as *FAR1*, another Zn(2)-Cys(6) transcription factor that regulates lipid metabolism related to appressorium turgor generation (Bin Yusof et al., 2014; Cruz-Mireles et al., 2024). RNA processing is another important area, with *CGRA01v4_06155* (*MGG_03116*) encoding a hypothetical protein involved in mRNA processing and splicing, homologous to the yeast *Prp19* (Bhadauria et al., 2010).

The study of effectors reveals that the genes *CGRA01v4_00409* (*GLRG_10780*) and *CGRA01v4_05892* (*GLRG_09920*) are predicted to be CFEM effectors CgCFEM19 and CgCFEM11, respectively, suggesting that they may play a role in pathogenicity. Although not functionally characterized, other CFEM effectors have been found to play a suppressive role in cell death activity during transient expression in *Nicotiana benthamiana* (Gong et al., 2020). *CGRA01v4_03968* (*MGG_13626*), a putative glycosyl hydrolase family effector, might induce cell death signaling in rice plants (Kim et al., 2013).
